# Evolution of taste processing shifts dietary preference

**DOI:** 10.1101/2024.10.11.617601

**Authors:** Enrico Bertolini, Daniel Münch, Justine Pascual, Noemi Sgammeglia, Carlos Ribeiro, Thomas O. Auer

**Author notes:** Corresponding author: Thomas O. Auer. equal contribution (Daniel Münch, Justine Pascual).

## Abstract

Food choice is an important driver of speciation and invasion of novel ecological niches. However, we know little about the mechanisms leading to changes in dietary preference. Here, we use the three closely-related species *Drosophila sechellia*, *D. simulans* and *D. melanogaster* to study taste circuit and food choice evolution. *D. sechellia,* a host specialist, feeds exclusively on a single fruit (*Morinda citrifolia*, noni) - the latter two are generalists living on various substrates. Using quantitative feeding assays, we recapitulate the preference for noni in *D. sechellia* and detect conserved sweet but altered bitter sensitivity via calcium imaging in peripheral taste neurons. Noni surprisingly activates bitter sensing neurons more strongly in *D. sechellia* due to a small deletion in one single gustatory receptor. Using volumetric calcium imaging in the ventral brain, we show that instead of peripheral physiology, species-specific processing of noni and sugar signals in sensorimotor circuits recapitulates differences in dietary preference. Our data support that peripheral receptor changes alone cannot explain altered food choice but rather modifications in how sensory information is transformed into feeding motor commands.

## Introduction

The decision to feed on or to reject a food source is fundamental for the survival of a species. While some animals feed on a broad variety of substrates, others are selective, specialised feeders that depend on the specific interaction with a single source of nutrients. These dietary shifts, often across closely-related species, have been associated with altered sensation of taste which is the initiating event in food consumption and main factor that motivates food intake or avoidance across all animals^1,2^. Taste cells express members of large families of chemosensory receptors, which show highly dynamic evolution^3,4^ with gains and losses across genomes. A well-known example is the giant panda, which lost a taste receptor sensing umami - the taste of protein - in its genome and lives exclusively on a bamboo diet^5^; cats lost their sense of sweetness through pseudogenisation of one of their sweet receptors^6^ and songbirds re-purposed their umami receptor for novel sweet detection^7^. In insects, dietary shifts often involve extreme specialisation on selected host plants. Plant cues from unripe citrus fruits and plant glucosinolates are detected by specifically tuned taste receptors in the Chinese citrus fly^8^ or *Pieris rapae*^9^, respectively. However, while these studies propose changes in individual receptors impacting taste cell physiology, a deeper understanding of signal processing after sensation at the periphery and its potential role in shaping food choice is missing.

Drosophilid flies, with the unique genetic toolset in *Drosophila melanogaster* (*D. melanogaster*) and closely-related species with diverse feeding habits offer a unique system to study this phenomenon. In *Drosophila*, like in mammals, gustatory sensory neurons (GSNs) localized on the fly’s main taste organ, the labellum of the proboscis, detect several taste modalities^2^, each of which promotes food acceptance or rejection^10,11^. Peripheral GSNs project their axons to the subesophageal zone (SEZ) of the fly brain^2^ where innervations are spatially separated by taste modality. Here, they connect to local modulatory interneurons and second-order neurons transmitting taste information either locally with reciprocal connectivity between neurons^12,13^ or to other brain areas^13–16^. The strict separation of taste modalities at the periphery quickly converges, already at second order neurons^13,17^, to eventually activate motor neurons driving feeding subprogrammes like proboscis extension or food ingestion^18^. This organization principle is conserved from flies to mammals^2,19^ but previous studies on taste circuit evolution mainly focused on the sensory periphery.

In the agricultural pest *D. suzukii*^20^, the herbivore fly *Scaptomyza flava*^21^ and the tropical island endemic *D. sechellia* living on a unique host fruit^22^, electrophysiological recordings of GSNs point towards shifts in sugar valuation (for *D. suzukii*)^20,23^, loss of bitter sensitivity (*all three*)^21,24,25^ or altered sweet/bitter integration (*D. sechellia*)^26,27^ impacting food choice. This altered peripheral physiology is likely due to gene losses and/or down-regulation of chemosensory genes for bitter^24,25,28,29^ or sweet detection^23^, including olfactory binding proteins^30^, gustatory^26,27^ or ionotropic receptors^26^. However, without functionally testing the role of these candidates in food preference in the respective species, evidence remains correlative and is mainly based on loss-of-function studies in *D. melanogaster*.

In addition, while these examples highlight the link of peripheral receptors and dietary preferences, how processing of taste information in downstream circuits evolves has not been investigated yet. Here, we study peripheral and central taste processing using the specialisation of *D. sechellia* on its selected host plant noni as an ethologically relevant paradigm for evolution of feeding preference.

## Results

### *D. sechellia* shows preference for noni in several feeding assays

To recapitulate the host preference of *D. sechellia* for noni in a semi-natural lab setting, we presented a group of flies four different fruits (banana, apple, grape, and noni) in a behavioural arena and filmed their spatial distribution over 24 hours. Consistent with the ecology of this species trio, *D. sechellia* shows a strong preference for noni while *D. simulans* and *D. melanogaster,* both generalist feeders that share a last common ancestor with *D. sechellia* 0.1-0.24 and 3-5 million years ago^22^, respectively, prefer the other fruits (**Fig. 1A**, respectively, prefer the other fruits**. 1A**). Importantly, this strong preference for noni in *D. sechellia* persists when testing near-anosmic flies (mutants for the two co-receptors Orco and Ir8a for antennal odorant and ionotropic receptors, respectively, **Fig. 1A, Sup. Fig. 1A**), which lose olfactory attraction to noni in an olfactory trap assay^31^. This indicates that other sensory modalities, apart from olfaction, contribute to the preference shift towards noni in *D. sechellia*.

**Figure 1.**
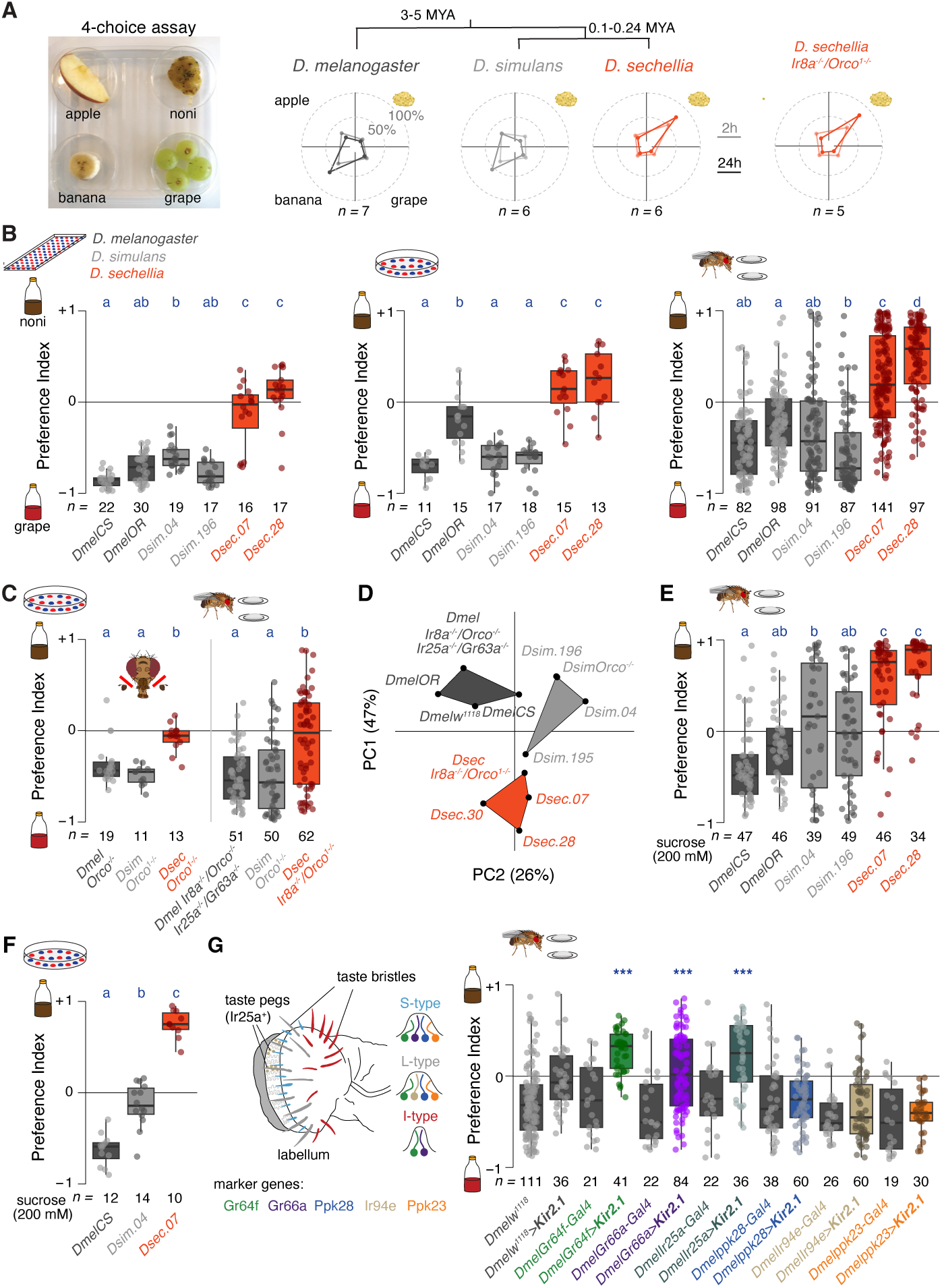
*D. sechellia* shifts preference from sugar to noni. **A)** Semi-natural 4-choice assay testing preference towards grape, banana, apple or noni fruit over 24 h. Spider plots: Percentage of flies (group of 20 males and 50 females) per quadrant 2 h (faded line) and 24 h (solid line) after assay start. Here and in other panels, sample sizes are indicated in the figure. *D.sechellia* = *Drosophila* Species Stock Center [DSSC] 14021-0248.07; *D. simulans* = DSSC 14021-0251.04; *D. melanogaster* = *CantonS*. On top: Phylogenetic relationship of the *D. melanogaster*, *D. simulans*, and *D. sechellia* trio. MYA = million years ago. Right: Distribution of near-anosmic *D. sechellia* (*Ir8a^-/-^/Orco^1-/-^*)^31^ flies at the same time points as shown for wild-type. **B)** Feeding preference for two strains of each species (*D.sechellia* = DSSC 14021-0248.07 and 14021-0248.28, *D. simulans* = DSSC 14021-0251.04 and DSSC 14021-0251.196, *D. melanogaster* = *CantonS* and *Oregon R*) for noni or grape juice in a 72-well (left), petri dish (middle) or flyPAD (right) two-choice assay. In the first two assays, preference index = (number of flies feeding on A -number of flies feeding on B)/number of total flies feeding. In the flyPAD, preference index = (number of sips on A – number of sips on B)/number of total sips. Similar phenotypes are observed for mutant strains lacking eye pigmentation (see **Sup. Fig. 1E**). For these and all other box plots, the center line represents the median, the box bounds the first and third quartiles, and the whiskers extend to the largest and smallest values within 1.5 x the interquartile range; individual data points are overlaid. For the 72-well and petri dish assays, a group of 80 or 20 males or females represents one datapoint, respectively. In the flyPAD, one individual female represents one datapoint. Kruskal-Wallis H test (72-well: H_(5)_=73.394, P=2.012e^-14^; petri dish: H_(5)_=64.468, P=1.445e^-12^; flyPAD: H_(5)_=238.45, P=2.2e^-16^). Pairwise comparisons were conducted using Dunn’s test with Bonferroni correction. For these and all other comparisons the results are shown in the figure, where groups sharing the same letter are not significantly different (P < 0.025). Exact p-values are listed in **Sup. Table 5**. **C)** Left: Feeding preference of Orco-mutant animals after antennal removal (rendering them anosmic) for noni vs. grape juice in the petri dish assay. Kruskal-Wallis H test (H_(2)_=19.227, P=6.681e^-05^; Dunn’s test results are shown in the figure). Right: feeding preference of olfactory mutant flies (*D. melangoaster Ir8a^-/-^/Orco^-/-^/Ir25a^-/-^/Gr63a^-/-^*, *D. simulans Orco^1-/-^*, *D. sechellia Ir8a^-/-^/Orco^1-/-^*) for noni vs. grape juice in the flyPAD. Kruskal-Wallis H test (H_(2)_=16.537, P=0.0002564; Dunn’s test results are shown in the figure). **D)** Principal component analysis of flyPAD data (**B,C, Sup. Fig. 2**) with clear species separation. *D.sechellia* = DSSC 14021-0248.07, 14021-0248.28, 14021-0248.30, *Ir8a^-/-^/Orco^1-/-^*; *D. simulans* = DSSC 14021-0251.04, DSSC 14021-0251.196, DSSC 14021-0251.195, *Orco^1-/-^; D. melanogaster* = *CantonS, OregonR, w^1118^*, *Ir8a^-/-^/Orco^-/-^/Ir25a^-/-^/Gr63a^-/-^*). **E)** Preference index for feeding on noni juice vs. 200 mM sucrose in the flyPAD. Kruskal-Wallis H test (H_(5)_=93.908 P=2.2e^-16^; Dunn’s test results are shown in the figure). **F)** Preference index for feeding on noni juice vs. 200 mM sucrose in the petri dish assay. Kruskal-Wallis H test (H_(2)_=28.526 P=6.393e^-07^; Dunn’s test results are shown in the figure). **G)** Left: schematic of the fly labellum depicting taste pegs and taste bristles. Below: Molecular markers for different neuron types in the short (S), intermediate (I) and long (L) sensilla. L- and S-sensilla house four, I-sensilla two neurons each. Right: Preference index for noni vs. grape juice feeding in control (*UAS-Kir2.1*) and experimental flies after silencing of selected taste cell populations. Kruskal-Wallis H test was performed among experimental and control flies for each taste cell population (Gr64f: H_(3)_=62.818, P=1.469e^-13^; Gr66a: H_(3)_=38.788, P=1.925e^-08^; Ir25a: H_(3)_=39.51, P=1.353e^-08^; ppk28: H_(3)_=15.84, P=0.001223; Ir94e: H_(3)_=24.31, P=2.152e^-05^; ppk23: H_(3)_=25.643, P=1.133e^-05^). Significant Dunn’s test results are shown in the figure (***P < 0.001) after Benjamini–Hochberg correction as pairwise comparison to Gal4-control.

We next analysed the role of taste in noni preference in several binary choice paradigms including a 72-well plate group assay^11^, a petri-dish group assay^32^ and the flyPAD (on single flies)^33^ using grape juice vs. noni juice as stimuli. The latter’s chemical composition is similar to noni fruit^31^ and it evokes comparable behavioural phenotypes^31,34^. In all three assays, *D. melanogaster* and *D. simulans* show a strong preference for grape juice. *D. sechellia* instead consumes more of the noni substrate than the other species (**Fig. 1B**) without significant sex differences (**Sup. Fig. 1B**) and with similar feeding rates across species (**Sup. Fig. 1C, D**). To test if olfactory attraction plays a role in the observed feeding phenotypes in these assays, we generated an *orco* allele in *D. simulans* (identical to *DsecOrco*^1^ in *D. sechellia*^31^, **Sup. Fig. 2A**). After removal of the antennae in Orco mutant flies of all three species (resulting in loss of all Or driven olfactory pathways; in the maxillary palp only Orco-dependent channels are present^35^), *D. simulans* and *D. melanogaster* still prefer grape juice in the petri dish assay. *D. sechellia* displays a neutral preference index, similar to results of olfactory impaired flies in the flyPAD assay (**Fig. 1C, Sup. Fig. 2B**). Principal component analysis of the flyPAD data (taking multiple parameters of feeding microstructure into account (**Sup. Fig. 2C, D**)) clearly separates *D. sechellia* from its two relatives, independently of having a functional olfactory system (**Fig. 1D**). These data support the view that, while olfaction contributes to noni attraction, taste is sufficient to establish species-specific food preference in this species trio.

Grape juice, grapes, banana and apples are highly enriched in sugars compared to noni juice and fruit^36^ (**Sup. Fig. 2E**). Contrary to *D. melanogaster* and *D. simulans*, which thrive on carbohydrate rich diets^36–38^, *D. sechellia* larvae exhibit developmental delays and show reduced lifespan as adults when fed on sugars, indicating a substantial shift in nutritional requirements. To test if sugar content is the main cause for diverse food preferences, we presented flies a choice between sucrose and noni juice in the flyPAD and petri dish assay (**Fig. 1E, F**). In this setup, *D. melanogaster* shows a strong preference for sucrose, *D. sechellia* almost exclusively feeds on noni while *D. simulans* displays an intermediate phenotype (in the flyPAD, **Fig. 1E**) or sugar preference (in the petri dish assay, **Fig. 1F, Sup. Fig. 2F**). Collectively, this analysis indicates that there is a switch in food consumption from grape and other, sugary fruits in *D. melanogaster* and *D. simulans* to noni in *D. sechellia* that can be attributed to the sugar preference differences across these species.

### Bitter and sweet sensation influence noni feeding in *D. melanogaster*

Flies use gustatory sensory neurons of their main taste organ, the labellum of the proboscis, to evaluate the suitability of a potential food source: Sweet sensing neurons located in taste bristles promote food consumption, bitter sensing neuron activation leads to an aversive response^1^ and taste peg neurons modulate feeding length^39^. Given the robust species-specific differences in noni vs. grape juice/sucrose consumption, we screened gustatory sensory neuron populations in *D. melanogaster* for their role in the observed species divergence. We selected Gal4 driver lines labelling all gustatory cell types in the labellum (**Fig. 1G**) and combined them with a *UAS-Kir2.1* transgene^40^ for targeted neuron silencing.

Compared to control genotypes, silencing sugar sensing gustatory bristle neurons using either *DmelGr64f-* or *DmeIlr25a-Gal4* (which is also expressed in taste pegs) led to a marked reduction in grape but not noni juice intake and a positive preference index for noni feeding (**Fig. 1G, Sup. Fig. 2G**). This is consistent with the hypothesis that the high sugar content of grape juice stimulates food intake through the activation of sweet sensing neurons. Silencing of the bitter sensing neuron population (using *DmelGr66a-Gal4*) markedly increased noni feeding, probably through a loss of aversive signals from this substrate, and experimental flies lost preference for one or the other food source (**Fig. 1G, Sup. Fig. 2G**).

The manipulation of ppk23 or ppk28 expressing neurons, which mark pheromone and water sensing cell populations^41,42^, respectively, did not lead to a significant change in preference but increased intake on any of the two substrates (**Fig. 1G, Sup. Fig. 2G**). Ir94e silencing, which is mainly involved in salt sensing^43^, did not alter intake nor preference. In sum, these results support a role of sweet and bitter sensing neurons, marked by Gr64f and Gr66a expression, and potentially taste pegs, in the decision of feeding on noni or grape substrate. Grape juice stimulates intake via activation of sweet sensing neurons while noni intake is reduced through activation of bitter sensing, aversive pathways in *D. melanogaster*.

### Comparison of sweet sensing circuits across species

Given that sweet and bitter sensing cells influence the decision to feed on noni or grape juice, we wanted to directly investigate these neuron populations in *D. sechellia.* Starting with sweet sensing neurons, we used CRISPR to integrate a Gal4 cassette into an early exon of *Ir25a* and *Gr64f* resulting in Gal4 reporters in the *cis*-regulatory environment of the native loci (**Sup. Fig. 3A**).

Reminiscent of expression in *D. melanogaster*, the *DsecIr25a^Gal4^*line is broadly expressed in the taste system, including taste pegs and bristles of the labellum, and olfactory sensory neurons^44–46^ (**Fig. 2A, Sup. Fig. 3B**). *DsecGr64f^Gal4^*expression is more restricted to sensory neurons in labellar taste bristles (**Fig. 2B, Sup. Fig. 3B, C**) and few olfactory sensory neuron populations as in *D. melanogaster*^10^. To be able to compare the anatomy of these circuits across species without confounding effects by the use of divergent transgenic tools, we generated homologous *Gr64f^Gal4^*-knock-in transgenics in *D. simulans* and *D. melanogaster*. Using these Gal4 reporters, we quantified the number of Gr64f expressing cells in labial palps, forelegs, midlegs, and hindlegs in all three species (**Fig. 2C, Sup. Fig. 3D, Sup. Table 1**). In contrast to a recent study in *D. suzukii*, which shows reduced sweet sensitivity and a remarkable one third reduction in *D. suzukii* labellar sweet sensing neurons^20^, we did not observe a strong species difference in Gr64f neuron numbers in the labellum or legs (**Fig. 2C, Sup. Fig. 3D**). This data indicates that loss of peripheral sugar sensing neurons does not explain the reduced sugar appetite of *D. sechellia*.

**Figure 2.**
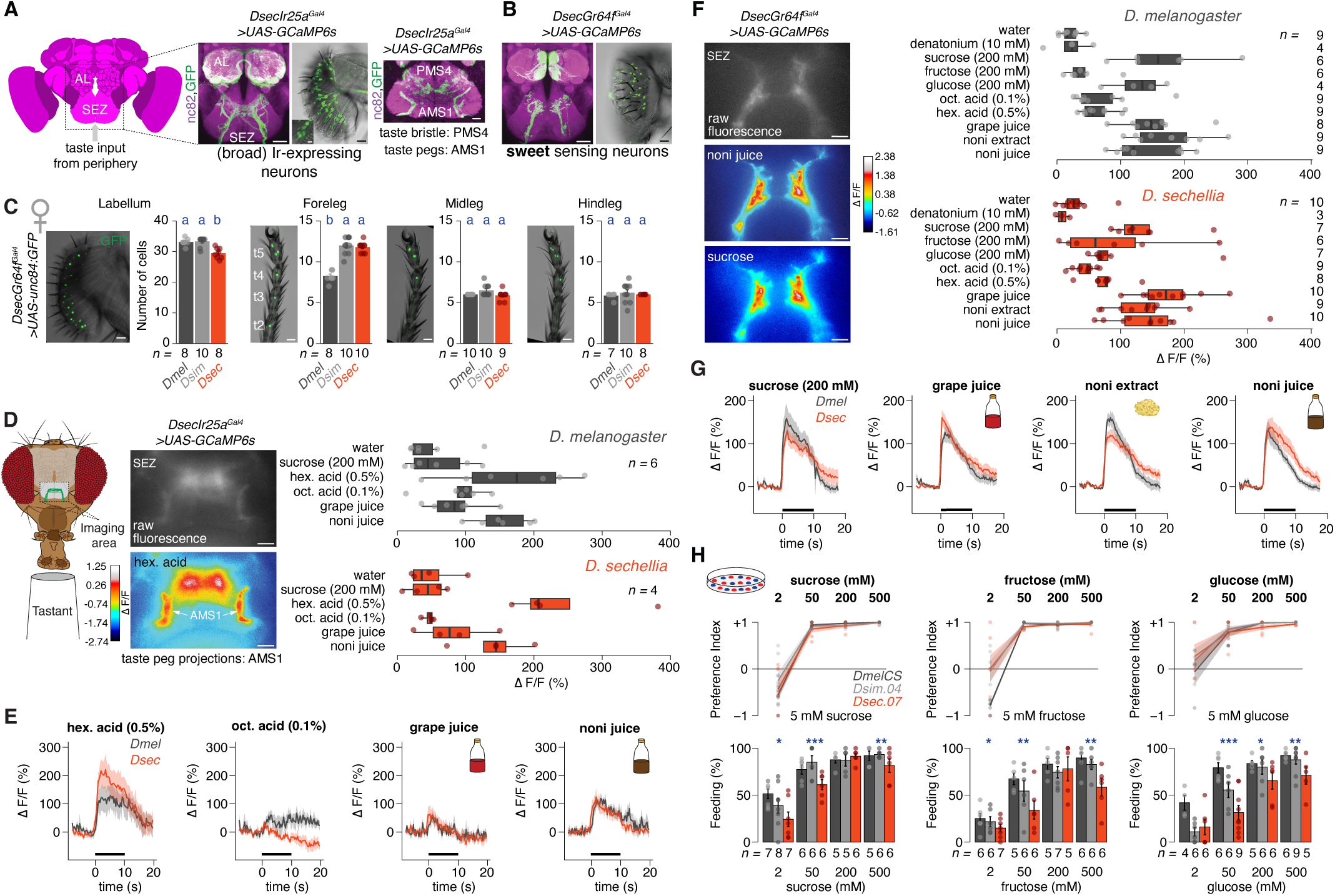
Taste peg and sweet neuron characterization. **A)** Left: schematic of the fly brain depicting the subesophageal zone (SEZ) receiving taste input from the periphery. AL = antennal lobe. Middle: Immunofluorescence with anti-GFP (recognizing GCaMP6s) and nc82 (labelling neuropils) antibodies in the brain of *D. sechellia Ir25a^Gal4^* transgenic flies expressing GCaMP6s. Scale bar = 25 µm. Next to the brain staining, the corresponding labelling in the labellum is shown (anti-GFP staining, scale bar = 25 µm). The inlet highlights taste pegs, scale bar = 5 µm. Right: Axonal innervation pattern in the SEZ. Axonal projections of taste bristle neurons target the PMS4 region; of taste peg neurons the AMS1 area. Scale bar = 10 µm. **B)** Immunofluorescence with anti-GFP and nc82 antibodies in the brain (left) or labellum (right, GFP only) of *D. sechellia Gr64f^Gal4^* transgenic flies expressing GCaMP6s. Scale bars = 25 µm. **C)** Number of Gr64f^+^ neurons in the labellum and legs of female *D. melanogaster*, *D. simulans* and *D. sechellia* based on transgenic labelling. In each panel, on the left a nuclear staining with an *UAS-unc84:GFP* reporter in the *D. sechellia* labellum and legs (scale bars = 25 µm). t5 – t2 = tarsal segments 5 – 2 (male cell numbers: **Sup. Fig. 2C, Sup. Table 1**). Here and in other panels, sample size is indicated in the figure. For these and all other cell number bar plots, the bar represents the mean, the error bar the standard error; individual data points are overlaid. Kruskal-Wallis H test (labellum: H_(2)_=14.05, P=0.0008894; foreleg: H_(2)_=17.631, P=0.0001484; midleg: H_(2)_=6.0282, P=0.04909; hindleg: H_(2)_=1.3862, P=0.5). Dunn’s test results (after Bonferroni correction) are shown in the figure. **D)** Left: Schematic of the calcium imaging set-up to characterize taste sensory neuron responses in the SEZ. Middle: Example image of raw fluorescence (top) and tastant evoked (0.5% hexanoic acid) calcium responses (bottom) in *D. sechellia* peg neurons (labeled by *Ir25a^Gal4^* driven GCaMP6s expression in the AMS1 area, scale bar = 10 µm). The colour scale depicts relative fluorescent changes (ΔF/F). Right: Quantification of tastant evoked calcium responses in Ir25a^+^taste peg neuron projections in the SEZ of *D. melanogaster* (top) and *D. sechellia* (bottom), reported as normalised GCaMP6s fluorescence changes. For these and all other physiology box plots, the centre line represents the median, the box bounds represent the first and third quartiles, and whiskers depict at maximum 1.5 x the interquartile range; individual data points are overlaid. Wilcoxon rank-sum test. No significant differences to *D. melanogaster* responses are detected. **E)** Temporal fluorescence changes in taste peg neuron projections during labellum stimulation. The solid line connects mean values of consecutive timepoints, the shaded area around the line defines the standard error of the mean. The horizontal black bar indicates the interval of tastant application. **F)** Left: Example image of raw fluorescence (top) and tastant evoked (noni juice) calcium responses (bottom) in *D. sechellia* sweet taste bristle neuron projections (labelled by *Gr64f^Gal4^* driven GCaMP6s expression, scale bar = 10 µm). The colour scale depicts relative fluorescent changes (ΔF/F). Right: Quantification of tastant evoked calcium responses in Gr64f^+^ taste bristle neuron projections in the SEZ of *D. melanogaster* and *D. sechellia*, reported as normalised GCaMP6s fluorescence changes. Wilcoxon rank-sum test. No significant differences to *D. melanogaster* responses are detected. **G)** Temporal fluorescence changes in sweet sensing taste bristle neuron projections during labellum stimulation. The solid line connects mean values of consecutive timepoints, the shaded area around the line defines the standard error of the mean. The horizontal black bar indicates the interval of tastant application. **H)** Top: Dose dependent feeding preference for individual sugars across species in the petri dish assay, 18-20 females per datapoint; sample size for each condition is indicated below. Bottom: percentace of flies feeding in the petri dish assay. Kruskal-Wallis H test preference index (5 mM sucrose H_(2)_=0.56183, P=0.7551; 50 mM sucrose H_(2)_=1.1057, P=0.5753; 200 mM sucrose H_(2)_=4.3406, P=0.1141; 500 mM sucrose H_(2)_=1.6403, P=0.4404; 5 mM fructose H_(2)_=2.2974, P=0.317; 50 mM fructose H_(2)_=4.0143, P=0.1344; 200 mM fructose H_(2)_=7.3313, P=0.2559; 500 mM fructose H_(2)_=1.3846, P=0.5004; 5 mM glucose H_(2)_=4.0495, P=0.132; 50 mM glucose H_(2)_=0.54317, P=0.7622; 200 mM glucose H_(2)_=4.783, P=0.09149; 500 mM glucose H_(2)_=0.5432, P=0.76). Kruskal-Wallis H test percentage of feeding (5 mM sucrose H_(2)_=8.6161, P=0.01346; 50 mM sucrose H_(2)_=15.888, P=0.0003549; 200 mM sucrose H_(2)_=0.73588, P=0.6922; 500 mM sucrose H_(2)_=11, P=0.004087; 5 mM fructose H_(2)_=6.4043, P=0.04068; 50 mM fructose H_(2)_=12.093, P=0.002366; 200 mM fructose H_(2)_=1.91, P=0.3848; 500 mM fructose H_(2)_=10.954, P=0.004182; 5 mM glucose H_(2)_=4.4752, P=0.1067; 50 mM glucose H_(2)_=18.076, P=0.0001188; 200 mM glucose H_(2)_=8.5812, P=0.0137; 500 mM glucose H_(2)_=11.642, P=0.002964).

Therefore, we turned to study the neurophysiology of taste pegs and sugar sensing taste bristle neurons. To this end, we combined our Gal4 drivers with *UAS-GCaMP6s* reporters in *D. sechellia* and *D. melanogaster* and stimulated the labellum with either noni or grape juice, selected noni acids^47^ or known agonists of sweet sensing pathways, while imaging calcium responses in the SEZ (**Fig. 2D**).

We first focused on taste pegs which are not accessible to electrophysiological recordings and have not been studied in *D. sechellia*. Taste pegs are known to respond to carbonation^45^, hexanoic acid^48^, and yeast stimuli and support feeding prolongation in *D. melanogaster*^39^. In both species, we detected a robust hexanoic acid response and noni juice led to increased neuronal activity in taste peg neuron projections compared to grape juice (**Fig. 2D, Sup. Fig. 3E** not normalised data). Overall, the activation pattern and temporal response to grape and noni juices in these neurons is similar between species (**Fig. 2E**) and indicates that physiological activity in peg neurons does not reflect *D. sechellia*’s specific preference for noni.

Next, we imaged calcium responses in Gr64f^+^ taste bristle neurons (**Fig. 2F**) which respond strongly to sucrose and glucose (**Fig. 2F, Sup. Fig. 3F**). Activation of these sweet sensing neurons is a strong driver of feeding initiation^1^ and their differential sensitivity could potentially explain why feeding preferences diverged between *D. melanogaster* and *D. sechellia*. However, we did not detect an increased sugar sensitivity in *D. melanogaster* compared to *D. sechellia* as suggested by *D. melanogaster*’s sucrose feeding preference. Moreover, when stimulated with noni or grape juice, *D. sechellia* Gr64f neurons responded stronger to grape than to noni juice (**Fig. 2F**) – contrary to the behavioural preference of *D. sechellia* for noni substrates. Taken together, responses in sweet sensing neurons, which show a similar temporal response profile across species (**Fig. 2G**), do not correlate with the divergent preference of *D. melanogaster* for sugars nor of *D. sechellia* for noni. Consistent with these physiological results, when presenting both species and *D. simulans* with the choice between a low and increasing concentrations of sugars (sucrose, fructose, glucose), we did not detect a behavioural difference in sugar sensitivity (**Fig. 2H**). This leads to the conclusion that the peripheral sweet sensing taste system in all three species does not significantly differ in its response to sweet or noni stimuli and therefore is not the main contributor to divergent food choices. Nevertheless, *D. sechellia* displayed a consistently reduced feeding rate on all three sugar substrates compared to its relatives (**Fig. 2H**). These data indicate that while all species have the same sensory discrimination ability for sugars, they differ in their ability to transform the sensory information into a sustained feeding motor pattern and hence sugar consumption.

### Altered neurophysiology in *D. sechellia* bitter sensing circuits

Previous studies and our neuron silencing experiments (**Fig. 1G**) propose altered bitter sensing as contributor to noni feeding across the *D. melanogaster/D. simulans/D. sechellia* trio^25–27^. This hypothesis is further supported by accelerated loss of (bitter sensing) gustatory receptors in *D. sechellia*^28,29^, the presence of several bitter substances like coumarin derivates in noni^49^ sensed by these receptors^50^, and the high acid content of this fruit potentially activating bitter sensing neurons^51^. To monitor neuronal activity in these circuits, we generated Gal4 reporter alleles at the *Gr66a* locus in all three species. As in *D. melanogaster*^11^ and *D. simulans*, the *D. sechellia* transgene drives expression in labellar taste bristle neurons projecting centrally to the SEZ (**Fig. 3A**).

**Figure 3.**
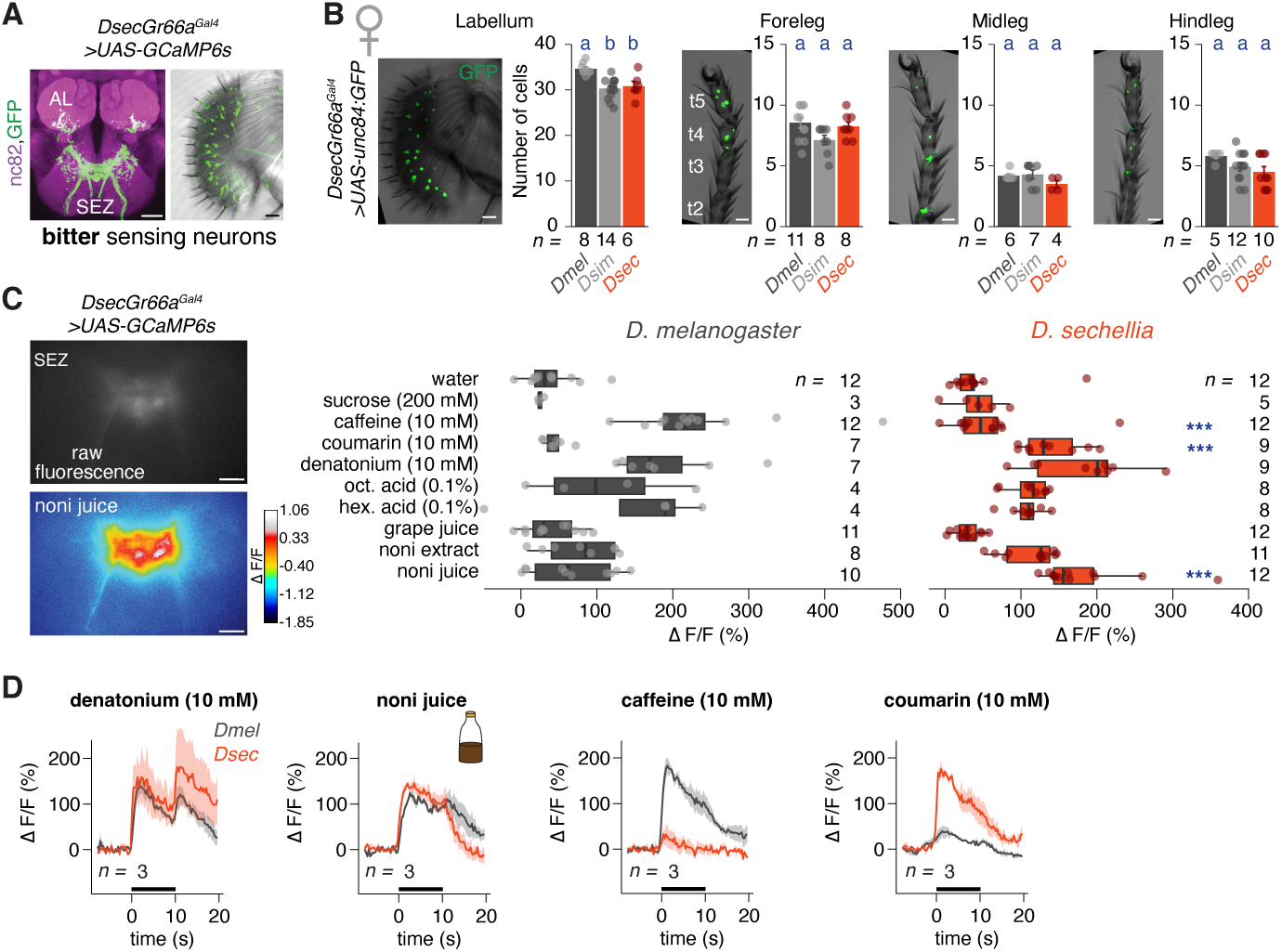
Bitter neuron numbers and physiology. **A)** Anti-GFP immunofluorescence in the SEZ and labellum of *DsecGr66a^Gal4^>UAS-GCaMP6s* transgenic flies. AL = antennal lobe. Scale bar = 25 µm. **B)** Number of Gr66a^+^ neurons in the labellum and legs of female *D. melanogaster*, *D. simulans* and *D. sechellia* based on transgenic labelling (males in **Sup. Fig. 4 A,** scale bar = 25 µm). Here and in other panels, sample size is indicated in the figure. For these and all other cell counts bar plots, the bar represents the mean, the error bar is the standard error; individual data points are overlaid. Kruskal-Wallis H test (labellum: H_(2)_=13.614, P=0.001106; foreleg: H_(2)_=4.8756, P=0.08735; midleg: H_(2)_=3.0446, P=0.2182; hindleg: H_(2)_=3.4719, P=0.1762). Dunn’s test results (after Bonferroni correction) are shown in the figure. **C)** Left: Example image of raw fluorescence (top) and tastant evoked (noni juice) calcium responses (bottom) in *D. sechellia* bitter taste bristle neuron projections (labelled by *DsecGr66a^Gal4^*>*GCaMP6s*, scale bar = 10 µm). The colour scale depicts relative fluorescent changes (ΔF/F). Right: Quantification of tastant evoked calcium responses in Gr66a+ bristles neuron projections of the fly labellum in the SEZ of *D. melanogaster* and *D. sechellia*, reported as normalised GCaMP6s fluorescence changes. Wilcoxon rank-sum test (***P < 0.001). Significant differences to *D. melanogaster* responses to the same stimulus are indicated in the figure (**Sup. Fig. 4C** for *D. simulans* data). **D)** Temporal fluorescence changes in bitter sensing taste bristle neuron projections during labellum stimulation. The solid line connects mean values of consecutive timepoints, the shaded area around the line defines the standard error of the mean. The horizontal black bar indicates the interval of tastant application. Shown is a subset of the whole Gr66a dataset as not all recordings could be temporally aligned.

We quantified the number of Gr66a^+^ neurons in the labellum and legs. Compared to previous work using Gr66a promoter-Gal4 transgenes^11,52^, we detected a higher number of Gr66a expressing cells in *D. melanogaster* using the knock-in allele (consistent with cell counts in electrophysiogical studies on labellar bitter neurons^11,25^). However, the number of bitter sensing neurons in *D. sechellia* was not reduced (apart from a slight reduction in the male labellum and hindlegs, **Fig. 3B, Sup. Fig. 4A**). This argues against dampened peripheral bitter inputs by altered Gr66a neuron counts in this species as observed e.g. in *D. suzukii*^24^.

Next, we characterized Gr66a circuit physiology in *D. melanogaster* and *D. sechellia* (**Fig. 3C**). Consistent with results from single sensillum recordings of labellar taste bristles^25^, we detected a marked reduction in caffeine sensitivity in *D. sechellia* whereas denatonium, hexanoic and octanoic acid responses were conserved (**Fig. 3C, Sup. Fig. 4B, C, D.** *simulans* and not normalised data). As in previous studies in *D. melangoaster*, we observed off-responses^53^ to a varying degree across stimuli (**Fig. 3D**) without an apparent species difference. Unexpectedly, *D. sechellia* bitter sensing neurons are highly sensitive to coumarin: instead of showing reduced sensitivity to this stimulant naturally found in noni fruit, coumarin leads to a stronger activation of Gr66a neurons in *D. sechellia* compared to their *D. melanogaster* homologs (**Fig. 3C**). Similarly, noni juice itself activates *D. sechellia* bitter sensing neurons more strongly, contradicting the hypothesis that loss of bitter/noni sensitivity drives the specialisation of *D. sechellia* to its noni host.

### Molecular determinants of differential responses in bitter neurons

The apparent loss of caffeine and gain of coumarin and noni sensitivity in *D.sechellia* bitter sensing neurons led us to investigate the genetic basis of this species difference. Accelerated loss of bitter receptors in *D. sechellia*^28,29^ has been proposed to change bitter neuron physiology and Gr33a, Gr39a, Gr66a and Gr93a were described as caffeine sensors in *D. melanogaster*^25,54,55^. Of these, the *Gr39a* locus is of particular interest. Transcripts from this locus are broadly expressed in bitter sensing neurons (**Fig. 4A**). They encompass four differentially spliced isoforms (*Gr39a.a-a.d*), and *Gr39a.b* (following FlyBase naming conventions) is pseudogenised in *D. sechellia*^28^ (8bp deletion, **Fig. 4B**). Moreover, this locus is evolutionary highly dynamic^56,57^ and potentially involved in altered taste physiology in herbivore drosophilids^21^. To test if pseudogenisation of *Gr39a.b* modifies caffeine and noni sensing in *D. sechellia*, we generated transgenic *D. sechellia* flies re-expressing the *DmelGr39a.b* receptor in Gr66a^+^ neurons. After labellar stimulation with caffeine, transgenic flies, however, did not show increased calcium activity in the SEZ compared to wild-type animals (**Sup. Fig. 5A**), arguing against a direct link between *Gr39a.b* pseudogenisation and loss of caffeine sensitivity in *D. sechellia*.

**Figure 4:**
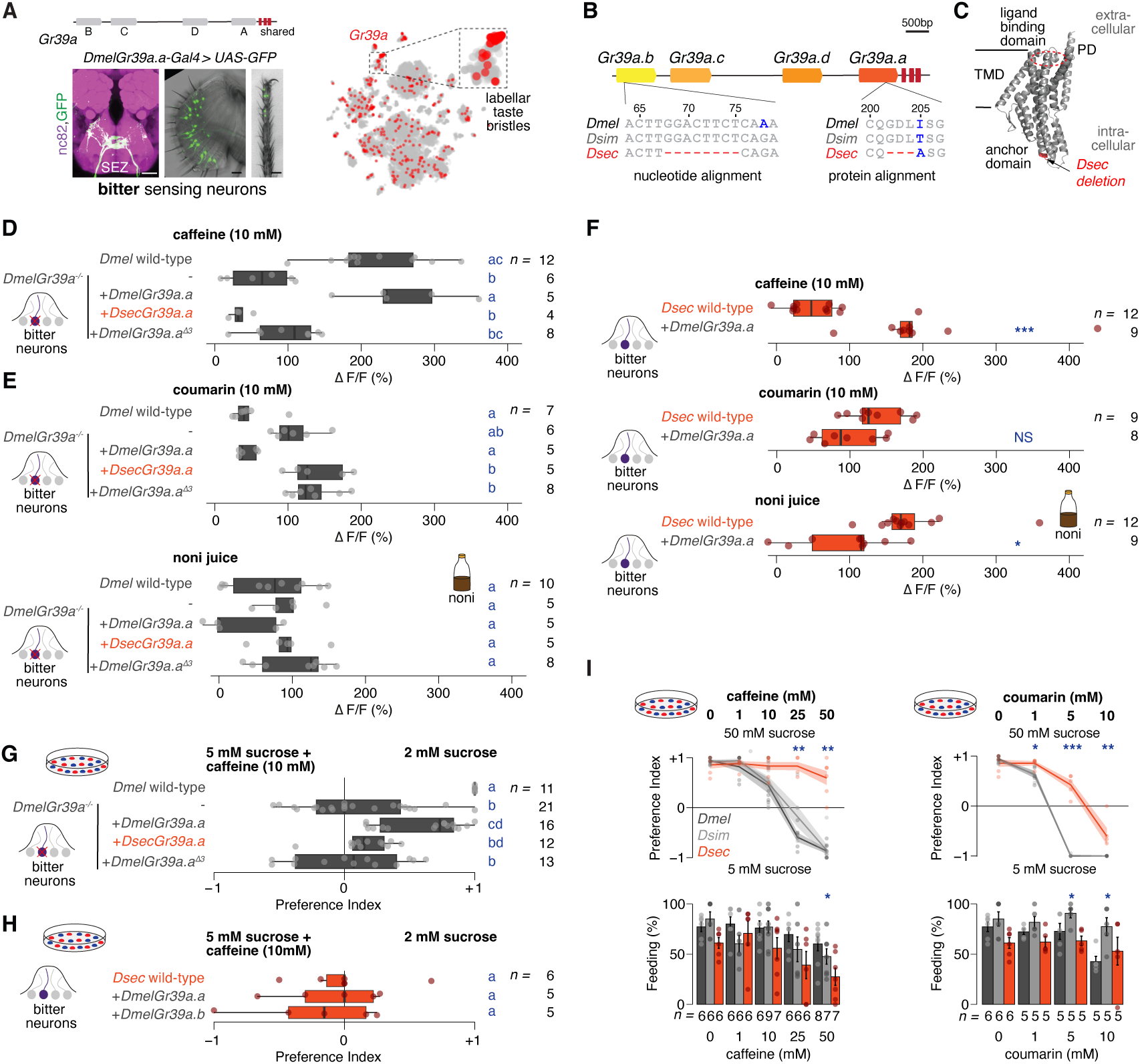
A small deletion in Gr39a.a recapitulates *D.sechellia* peripheral sensory neuron physiology. **A)** Top: Schematics of the *Gr39a* locus with four splice isoforms (grey boxes) sharing the last three common exons (red boxes). Below: Anti-GFP (and nc82) immunofluorescence in the SEZ, labellum and foreleg of *D. melanogaster Gr39a.a-Gal4 >UAS-GCaMP6s* transgenic flies. Scale bars = 25 µm. Right: Expression of Gr39a (in red) in individual neurons of the fly proboscis and maxillary palp (data from the FlyCellAtlas^85^). **B)** Of the four different transcripts of the Gr39a locus (*Gr39a.a*, *Gr39a.b*, *Gr39a.c*, *Gr39a.d*) a 8bp deletion in the most distal isoform (here called *Gr39a.b* following the Flybase nomenclature; in some studies *Gr39a.a*^56^), shown in a nucleotide alignment of all three species, leads to a frameshift and pre-mature stop codon after 33 residues in *D. sechellia*. Numbers indicate nucleotide position in the open reading frame. Right: The *Gr39a.a* protein carries a *D. sechellia*-specific three amino acid deletion (residues 202-204). Shown here a protein alignment of all three species. Numbers indicate protein residue number. In blue: divergent nucleotides and residues. **C)** Predicted protein structure of *D. melanogaster* Gr39a.a (monomer). The ligand binding domain faces the extracellular space next to the ion-conducting pore domain (PD). The intracellular anchor domain is important for tetramer assembly and the three amino acid deletion in *D. sechellia* is near this domain. TMD = transmembrane domain. The domain annotation is based on Ref.^59–62^. **D)** Quantification of caffeine evoked calcium responses in Gr39a.a^+^ bristle neuron projections of the fly labellum in the SEZ of *D. melanogaster* in wild-type (top) and *Gr39a^-/-^* mutant animals re-expressing either the *D. melanogaster* (*DmelGr39a.a*), *D. sechellia* (*DsecGr39a.a*) or *D. melanogaster* orthologue carrying the *D. sechellia* specific 3 amino acid deletion (*DmelGr39a.a ^Δ3^*). Here and in other panels, sample size is indicated in the figure. Kruskal-Wallis H test (H_(4)_=26.343, P=2.698e^-05^). Dunn’s test results (after Bonferroni correction) are shown in the figure. **E)** Quantification of coumarin (top) and noni juice (bottom) evoked calcium responses in Gr39a.a^+^ bristle neuron projections of the fly labellum in the SEZ of *D. melanogaster* of the indicated genotypes. Kruskal-Wallis H test (coumarin: H_(4)_=20.899, P=0.0003316 ; noni juice: H_(4)_=6.5233, P=0.1633). Dunn’s test results (after Bonferroni correction) are shown in the figure. **F)** Quantification of caffeine (top), coumarin (middle), and noni juice (bottom) evoked calcium responses in Gr66a^+^ bristle neuron projections of the fly labellum in the SEZ of *D. sechellia* in wild-type and animals expressing *D. melanogaster Gr39a.a* (*DmelGr39a.a*). Wilcoxon rank-sum test (caffeine: W = 0.29167, p = 0.5892; coumarin: W = 3, p = 0.08326; noni juice: W = 8.9423, p = 0.01); (***P < 0.001; **P < 0.01). Dunn’s test results (after Bonferroni correction) are shown in the figure. **G)** Preference index for caffeine (10 mM) in 5 mM sucrose vs. 2 mM sucrose feeding in the petri dish assay for *D. melanogaster* of the indicated genotypes. Kruskal-Wallis H test (H_(4)_=31.118, P=2.896e^-06^). Dunn’s test results (after Bonferroni correction) are shown in the figure. **H)** Preference index for caffeine in 5 mM sucrose vs. 2 mM sucrose feeding in the petri dish assay for *D. sechellia* of the indicated genotypes. Kruskal-Wallis H test (H_(2)_=0.26381, P=0.8764). Dunn’s test results (after Bonferroni correction) are shown in the figure. **I)** Top: Preference index for feeding on increasing concentration of caffeine (left) or coumarin (right) in 50 mM sucrose vs. 5 mM sucrose in the petri dish assay for all three species. 18-20 females per datapoint; sample size for each condition is indicated below. Bottom: Percentage of flies feeding in the petri-dish assay. Kruskal-Wallis H test preference index (1 mM caffeine H_(2)_=1.2832, P=0.5264; 10 mM caffeine H_(2)_=5.1312, P=0.07687; 25 mM caffeine H_(2)_=9.291, P=0.009605; 50 mM caffeine H_(2)_=13.005, P=0.0015; 1 mM coumarin H_(2)_=6.719, P=0.03475; 5 mM coumarin H_(2)_=12.118, P=0.002336; 10 mM coumarin H_(2)_=9.8824, P=0.007146). Kruskal-Wallis H test percentage of feeding (1 mM caffeine H_(2)_=2.8511, P=0.2404; 10 mM caffeine H_(2)_=2.9132, P=0.233; 25 mM caffeine H_(2)_=2.8657, P=0.2386; 50 mM caffeine H_(2)_=7.4973, P=0.02355; 1 mM coumarin H_(2)_=4.4585, P=0.1076; 5 mM coumarin H_(2)_=6.5215, P=0.03836; 10 mM coumarin H_(2)_=6.0304, P=0.04904). (***P < 0.001; **P < 0.01; *P < 0.025).

Focusing on the other splice isoforms, we detected a *D. sechellia*-specific deletion of three amino acids (Δ202-204) in Gr39a.a (**Fig. 4B**). Modelling of Gr39a.a with AlphaFold^58^ and comparison to recently published structures of insect gustatory sweet receptors^59–62^ suggest that these residues fall within the intracellular domain of the receptor close to the anchor domain. This domain is involved in tetramer assembly of receptor subunits but not ligand binding (**Fig. 4C**). We tested the functional consequence of this three amino acid deletion first in *D. melanogaster* flies. To this purpose, we generated transgenic flies expressing either *Dmel*Gr39a.a or *Dsec*Gr39a.a in Gr39a.a^+^ neurons lacking transcripts from the endogenous locus (*Gr39a^-/-^*). When presenting caffeine to the labellum of *Gr39a^-/-^* mutants, they completely lack the usually strong physiological response of wild-type individuals (**Fig. 4D**). Re-expression of *Dmel*Gr39a.a fully rescues physiological caffeine responses while the *Dsec*Gr39a.a orthologue does not restore caffeine sensitivity. To control for the effect of epistatic interactions with other residues of the receptor, we deleted the three amino acids absent in *D. sechellia* in the *D. melanogaster* Gr39a.a receptor. Testing this *Dmel*Gr39a.a^Δ3^ isoform resulted in similar phenotypes as the *D. sechellia* orthologue, supporting that this deletion leads to a failure of caffeine detection by this receptor (**Fig. 4D**). In sum, these experiments show that a three amino acid deletion in *Dmel*Gr39a.a recapitulates the *D. sechellia* Gr39a.a receptor phenotype and caffeine responses detected in endogenous *D. sechellia* bitter neurons.

As we observed additional physiological alterations in Gr66a^+^ neurons in *D. sechellia*, namely increased responses to noni and coumarin, we wanted to clarify if these were affected by the Gr39a locus. To this end, we presented to the same *D. melanogaster* transgenics introduced above coumarin and noni juice (**Fig. 4E**). Interestingly, while *D. melanogaster* wild-type flies do not respond strongly to any of the two, sensitivity to coumarin (and not significantly to noni) is increased upon loss of the *Gr39a locus*. Consistent with the caffeine rescue results, re-expression of *Dmel*Gr39a.a restores a wild-type phenotype. Re-expression of *Dsec*Gr39a.a or *Dmel*Gr39a.a^Δ3^ instead recapitulates the physiology of *Gr39a^-/-^* mutants (**Fig. 4E**). These results indicate that loss of caffeine and gain of coumarin and noni sensitivity are caused by the same genetic modification.

To clarify, if introduction of *Dmel*Gr39a.a has an impact on neuron physiology in *D. sechellia*, we expressed this receptor in endogenous Gr66a+ neurons in this species. While wild-type *D. sechellia* flies’ Gr66a neurons do physiologically not respond to caffeine stimulation, these “rescue” animals gained caffeine sensitivity (**Fig. 4F**). Consistent with a concurrent alteration in coumarin and noni sensitivity, responses to both stimuli in *D. sechellia* Gr66a neurons were reduced by expression of the *Dmel*Gr39a.a receptor compared to wild-type flies (**Fig. 4F**). Taken together, these experiments suggest that sequence changes in the Gr39a.a receptor alone can explain the physiological differences towards several tastants observed in *D. melanogaster* and *D. sechellia* endogenous bitter sensing neurons. In *D. melanogaster*, a functional Gr39a.a receptor confers sensitivity to caffeine and dampens responses to coumarin and noni juice. In *D. sechellia*, a three amino acid deletion most probably renders *Dsec*Gr39a.a non-functional resulting in a loss of caffeine responses and gain of coumarin and noni sensitivity in Gr66a neurons.

Given that Gr66a neuron activity is a good predictor of behavioural aversion to a selected stimulus^1,11^, we next tested the behavioural impact of Gr39a.a isoform expression in *D. melanogaster*. In a two-choice petri dish assay, *D. melanogaster* wild-type flies show complete aversion towards 5 mM sucrose with added caffeine compared to 2 mM sucrose (**Fig. 4G, I**). Consistent with the Gr66a neuron physiology described above, loss of the *Gr39a* locus significantly reduces aversion to a neutral preference index which is rescued to (almost) wild-type levels upon re-expression of *Dmel*Gr39a.a. Re-expression of *Dsec*Gr39a.a instead fails to rescue this aversive behaviour, as does *Dmel*Gr39a.a^Δ3^ (**Fig. 4G**) indicating that peripheral Gr66a neuron responses are directly translated into feeding preference in *D. melanogaster*. Similarly, *D. sechellia* wild-type flies, which lack peripheral caffeine responses, showed an increased tendency to feed on caffeine containing food when compared to *D. melanogaster* (**Fig. 4H, I**)^25,27^. However, contrary to the results in *D. melanogaster*, gain of physiological caffeine responses through re-expression of *Dmel*Gr39a.a does not result in behavioural aversion towards caffeine in this species (**Fig. 4H**). This indicates that peripheral sensing of bitter tastants is processed differently in *D. sechellia* downstream circuits.

To clarify if this phenomenon is specific for caffeine, we tested food consumption of wild-type flies in the two-choice petri dish assay towards increasing concentrations of caffeine and coumarin in 50 mM sucrose vs. 5 mM sucrose. *D. sechellia* display a reduced sensitivity to caffeine matching the bitter neuron physiological data. Coumarin instead, which strongly activates *D. sechellia* bitter neurons, does not induce strong behavioural aversion to the same degree as in *D. melanogaster* and *D. simulans*^27^ (**Fig. 4I**) similar to the *Dmel*Gr39a.a rescue results. Nevertheless, behavioural bitter aversion is not generally impaired in *D. sechellia* as denatonium avoidance is conserved across species (**Sup. Fig. 5B**). Collectively, these results indicate that while for some compounds like caffeine and denatonium, the physiology in endogenous bitter neurons predicts aversive feeding behaviours, for others like coumarin and noni, activation of bitter neurons does not lead to behavioural aversion in *D. sechellia*. This led us to conclude that, similarly to sugar, the processing of peripheral bitter information is tastant specific and altered in *D. sechellia* compared to *D. melanogaster*.

### Altered sensorimotor processing reflects behavioural diversification

Taste information from the labellum is transferred from peripheral sensory neurons over several layers of interneurons to motor neurons (MNs)^18^. These initiate feeding via proboscis extension or food ingestion through proboscis pumping after activation of muscles in the fly proboscis (**Fig 5A**)^14^. Recent circuit reconstructions have delineated the circuitry leading to proboscis extension from the sensory periphery to motor neuron 9^15^ and motor neurons arborizing in the superficial neuropil of the SEZ control pharynx pumping (motor neurons 10-13)^14^. To investigate how evolution may have modified the transformation of sensory information into motor outputs along the sensory-motor arc, we leveraged the power of volumetric imaging to capture activity across the entire SEZ^18^. We hypothesised that this comprehensive approach should reveal differences in tastant-evoked responses in higher-order taste circuits of *D. melanogaster*, *D. simulans*, and *D. sechellia*, providing insights into how sensory signals are processed and relayed to motor pathways.

**Figure 5.**
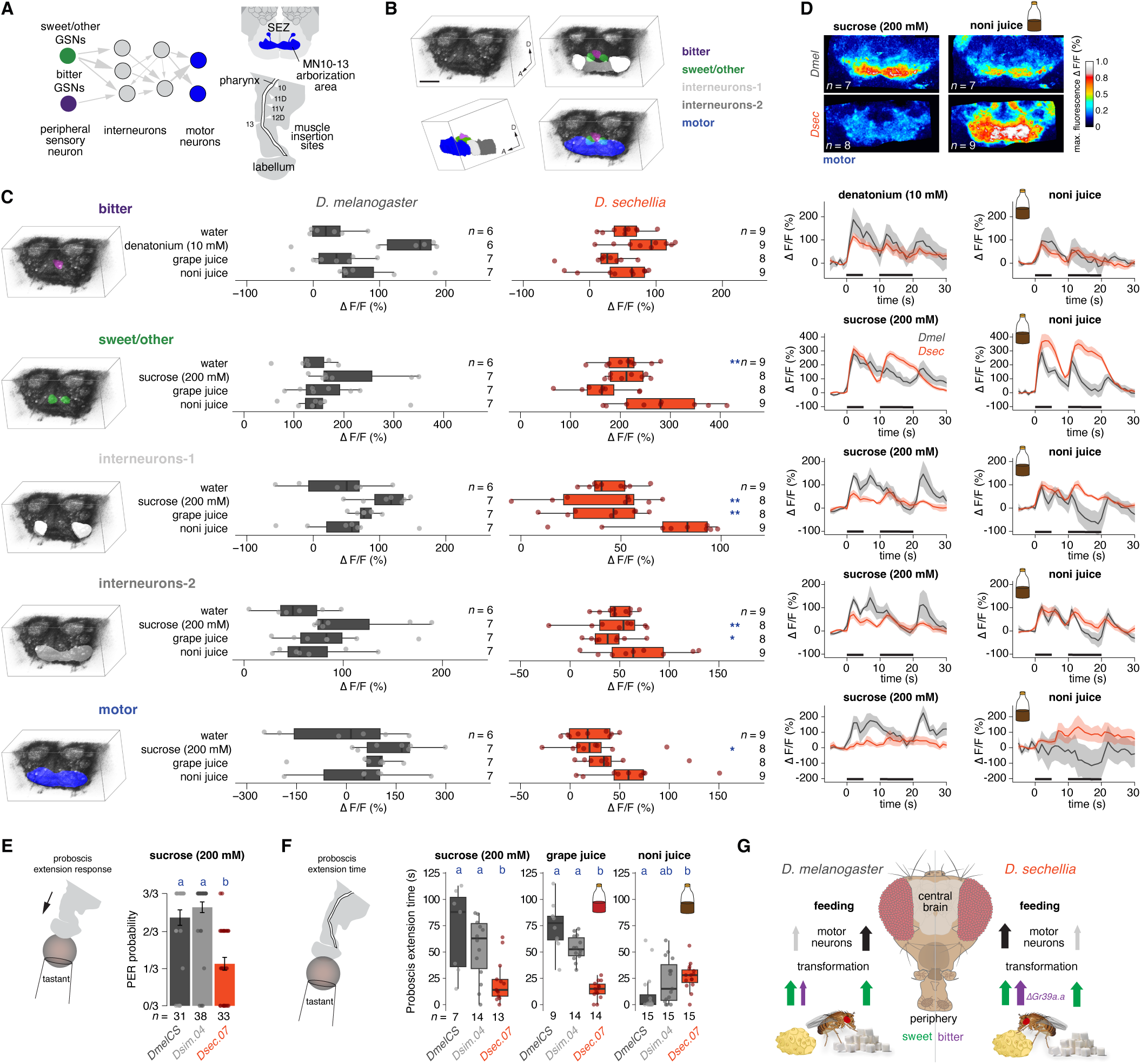
Volumetric imaging in the ventral brain uncovers processing differences between species. **A)** Left: Schematics of simplified peripheral, interneuron and motor neuron circuitry in the SEZ. Arrows indicate synaptic connections. Right: schematic arborization area of motor neurons (MN) 10-13 in the SEZ. Below: schematics of the fly pharynx in the proboscis with muscle insertion sites (white arrowheads) of muscles innervated by motor neurons 10-13. Based on data from Ref^14^. **B)** Top left: Imaging volume in the *D. sechellia* SEZ (anterior view, fluorescent signal inverted; A, anterior; D, dorsal). Right, top: Segmentation of bitter sensing projections (purple), sensory sweet/other sensing projections (green) and interneuron regions 1 and 2 (different shades of grey). Bottom: Segmentation of bitter sensing projections (purple), sensory sweet/other sensing projections (green) and motor neuron arborizations (blue). Bottom left: Lateral view of segmented regions. **C)** Left: Segmentation of imaging volume. Middle: Quantification of tastant evoked neuronal activity (maximum responses) in the segmented regions in *D. melanogaster* and *D. sechellia*. Here and in other panels, sample size is indicated in the figure. Wilcoxon rank-sum test (**P < 0.01; *P < 0.05). Significant differences to *D. melanogaster* responses are shown in the figure. For visualization purposes, one outlier (*D. melanogaster/*motor region/noni juice ΔF/F= −438%) is excluded from the display but included in the statistical analysis. Right: Temporal fluorescence changes in segmented areas during labellum stimulation. The solid line connects mean values of consecutive timepoints, the shaded area around the line defines the standard error of the mean. The horizontal black bar indicates the interval of tastant application. See **Sup. Figure 5** for *D. simulans* data. **D)** Maximum intensity projections of fluorescent signals in the motor region upon presentation of sucrose or noni juice in *D. melanogaster* (top) and *D. sechellia* (bottom). ΔF/F volumes of each recording were averaged across animals, and maximum intensity projections were applied in the time dimension to derive the maximum response per voxel. Sample size is indicated in the figure. **E)** Probability of proboscis extension response towards 200 mM sucrose in *D. melanogaster*, *D. simulans*, and *D. sechellia* in three consecutive trials. Kruskal-Wallis H test (H_(2)_=39.073, P=3.276e^-09^). Dunn’s test results (after Bonferroni correction) are shown in the figure. **F)** Time of proboscis extension in three species when presented with sucrose (200 mM), grape or noni juice. Kruskal-Wallis H test (sucrose: H_(2)_=11.173, P=0.003748; grape juice: H_(2)_=28.491, P=6.504e^-07^; noni juice: H_(2)_=9.0659, P=0.01075). Dunn’s test results (after Bonferroni correction) are shown in the figure. **G)** Schematics depicting food preference and taste information processing in *D. melanogaster* and *D. sechellia*. Peripheral signals of sweet and bitter sensing circuits are transmitted to the central brain. Sugars activate motor neurons for feeding initiation and ingestion in *D. melanogaster*. In *D. sechellia*, altered transformation of peripheral sugar and noni signals lead to motor neuron activation mainly by noni.

We expressed GCaMP6s pan-neuronally^31,63^ and imaged neuronal calcium signals across the whole SEZ upon stimulation with noni, grape juice, denatonium, and sucrose. For each species, we generated an average brain response and segmented functional areas based on anatomical and functional landmarks. Following this approach, we could distinguish five regions in all three species: the first region contains taste bristle innervations of bitter sensing neurons from the sensory periphery (bitter); the second innervations of peripheral sweet/other sensing populations (sweet/other). Flanking these sensory input zone, we identified two brain regions containing mainly interneurons (interneurons-1 and interneurons-2) and lastly one region representing motor neuron outputs (motor, **Fig. 5B**).

In the region mainly innervated by projections of bitter sensing neurons (bitter), *D. sechellia* did not show a reduced noni response compared to *D. melanogaster* and *D. simulans* (but rather an increase compared to the latter) confirming our results with the *Gr66a^Gal4^*driver (**Fig. 5C**). In the neighbouring regions receiving input from peripheral sweet (Gr64f^+^), water, and other sensory neuron types (sweet/other), we detected an increase in water and noni responses in *D. sechellia* (**Fig. 5C**). Interestingly, this region is also strongly responding to denatonium (**Sup. Fig. 5C**) supporting the heterogeneity of innervations in this area, including bitter neurons. Next, we compared the activity in two regions downstream of peripheral sensory neurons, containing most probably SEZ interneurons involved in taste signal processing (interneurons-1 and interneurons-2). Consistent with the feeding behaviour of all three species, these regions show the highest neuronal activity when sucrose is presented to the labellum in *D. melanogaster* (and to a lesser degree in *D. simulans*, **Sup. Fig. 5D**). Importantly, both regions respond most strongly to noni juice in *D. sechellia* (**Fig. 5C**). Moreover, sucrose fails to induce a significant response in these regions in *D. sechellia* supporting the idea that the strong sugar response of *D. melanogaster* taste downstream circuits^13^ is lost in this species. Similarly, in the motor neuron arborization region of MNs10-13 in the anterior part of the SEZ (motor), *D. sechellia* lacks a significant sucrose but shows an increased noni response (**Fig. 5C, D**; albeit not significantly different compared to *D. melanogaster*). *D. simulans* displays an intermediate phenotype (**Sup. Fig. 5D**), reflecting its consumption behaviour (noni vs. scurose) in the flyPAD (**Fig. 1E**). Overall, these results support the idea that species differ in how they transform peripheral taste information within higher order sensorimotor circuits. While *D. melanogaster* (and *D. simulans*) maintain a strong sucrose response from the periphery to motor outputs, in *D. sechellia* noni, but not peripheral sugar responses are efficiently propagated to feeding motor neurons and hence promote feeding.

The neuronal activity patterns in the SEZ of all three species nicely match their behaviour when presented with noni vs. grape juice or sucrose in the two-choice feeding assays (**Fig. 1**). Nevertheless, as these assays score food consumption, we aimed to establish a more direct link between peripheral taste sensation and motor neuron output. To this purpose, we presented sucrose to the labellum of *D. melanogaster*, *D. simulans* and *D. sechellia* and measured the probability of proboscis extension response (PER, **Fig. 5E**) – a motor programme necessary to initiate feeding. Reflecting the loss of sucrose signal propagation to motor neurons in *D. sechellia*, these flies show a significant reduction in PER probability upon sucrose presentation supporting the reduced potential of sugars to promote consumption. To monitor the vigor of food intake (reflecting activity of MNs10-13), we measured the duration of proboscis extension as a proxy for the rate of feeding upon presentation of sucrose, grape, or noni juice (**Fig. 5F**). We offered individual tastants to immobilized flies and summed up the time the proboscis remained extended across species upon stimulus encounter^64^ (**Fig. 5F**). In support of the strong feeding preference for sugar in *D. melanogaster* and *D. simulans* and reflecting the activity in pumping related motor regions, we observed long proboscis extension times when offering sucrose or grape juice to both species. In contrast, *D. sechellia*’s proboscis extension time was very low for both stimuli. Presentation of noni instead significantly increased proboscis extension time in contrast to the other two species (**Fig. 5F**).

In sum, these experiments align with our pan-neuronal imaging data suggesting that, despite similar peripheral taste neuron responses, sugar substrates lead to higher motor neuron activity and subsequently feeding in *D. melanogaster* compared to the noni specialist *D. sechellia* which lost sugar signal propagation and appetite.

## Discussion

We used the specialisation of *D. sechellia* on its selected host fruit noni as a paradigm to study taste and feeding evolution in insects (**Fig. 5G**). Despite the strong olfactory attraction of *D. sechellia* to noni, our and others’ experiments in smell-blind flies^26,27^ support that olfaction alone is not sufficient to explain the divergent preference for this fruit across the *D. sechellia*/*D. simulans*/*D. melanogaster* species trio. While it is highly likely that multi-modal integration of olfactory and gustatory cues happens in the central brain^65^, our physiological data, performed on animals lacking antenna and maxillary palps, clearly shows differential taste related signals in the SEZ and beyond.

Our calcium imaging analysis go beyond previous attempts to characterize the taste system in *D. sechellia* using single sensillum recording as we monitor the population response of all neurons expressing our Gal4 drivers at the same time. In support of the previously reported electrophysiological loss of caffeine and bitter responses^25^, we can recapitulate this trait in our settings. Similarly, we detect comparable attractive responses in *D. sechellia* and *D. melanogaster* towards the two noni-enriched acids octanoic and hexanoic acid^26,66^. However, while differences in sweet and acid integration between *D. melanogaster* and *D. sechellia* might happen already at the level of peripheral taste neurons^26,66^, this does not have a major impact on noni sensing. Contrary to previous hypotheses, we detected a surprising increase in bitter neuron responses towards noni in *D. sechellia* which we could ascribe to an individual short deletion in the *Gr39a.a* gene. Our data indicate that the *D. sechellia* specific deletion in the Gr39a.a paralog -instead of the pseudogenisation of *Gr39a.b* - renders the receptor non-functional through a (hypothesised) failure to assemble a functional receptor complex. The resulting altered physiology in *D. sechellia* supports that interactions of Grs (which probably form heteromeric tetramers) are complex and can lead to inhibitory, non-linear interactions amongst receptors^11,25^. Apart from mutations in ligand binding sites, which are evolutionary hotspots in olfactory receptor genes^31,67^, Gr receptor-receptor interaction domains could represent an alternative target to modulate gustatory responses in neurons expressing multiple Grs by altering receptor-complex assembly and function.

The reduced caffeine sensitivity in peripheral neurons and the resulting loss of behavioural aversion in *D. sechellia* is in line with previous studies linking sensory neuron responses with behaviours in drosophilids^20,21,25–27^. However, the increased sensitivity to noni in *D. sechellia* hints towards altered processing or integration of bitter signals in (local) interneurons in the SEZ and beyond^13,17–19^. These circuits, or modulatory systems influencing sweet and bitter integration^68–71^, might have undergone evolutionary re-modelling to block a strong aversive response towards noni in *D. sechellia*.

Indeed, our volumetric imaging experiments indicate that signal processing in downstream inter- and motor-neurons is altered across species. While peripheral sweet sensing neuron responses are conserved, sugars do not activate motor regions to the same degree in *D. sechellia*. The downstream circuitry of sugar sensing neurons has been mapped in *D. melanogaster* in great detail^13^ from the periphery to motor output. Our data suggest that along this sensorimotor processing axis, the high efficiency by which sugar taste signals can recruit feeding motor neurons, characteristic for *D. melanogaster* (and *D. simulans*), is lost in *D. sechellia*. This data is reminiscent of results in *D. suzukii*, where changes in the processing of conserved peripheral sugar sensitivity leads to divergence behavioural output (in this case in oviposition) across species^20,23^.

The reduced feeding rate and loss of motor region activation in *D. sechellia* by sucrose is in line with additional changes happening in ingestion circuits in this species to match its altered nutritional landscape. *D. sechellia*’s lost ability to metabolize high sugar diets and the negative effects of a carbohydrate rich environment on this species^36–38^ prevent high consumption of sweet foods. Future comparative studies with *D. melanogaster* will show how other carbohydrate mediated signalling pathways e.g. in the brain^72^, gut-brain axis^73^ and reproductive system^74^ are affected by the dietary change of *D. sechellia* and which mechanisms exist to reach nutrient homeostasis across different feeding regimes.

## Author contribution

T.O.A. and E.B. conceived the project. All authors contributed to experimental design, analysis and interpretation of results. Specific experimental contributions were as follows: E.B. performed molecular cloning, behavioural assays, immunostainings, and physiological experiments. D.M. performed volumetric calcium imaging. J.P. performed molecular cloning, behavioural assays, and immunostainings. N.S. performed behavioural assays. C.R. provided guidance to D.M. T.O.A. performed molecular cloning, immunostainings, preliminary physiological experiments, and generated transgenic lines. T.O.A. and E.B. wrote the paper with input from all authors. All authors read and approved the final version of the manuscript.

## Acknowledgements

We thank H. Dweck, B. Prud’homme, J. Simpson, H. Dionne, G. Rubin, S. Caron, and J. M. Bateman for sharing fly strains and reagents, N. Himmel for help in protein structure prediction, F. Meyenhofer for help in image quantification automation and A. Rebelo Pimentel for help with embryo alignments for microinjections. We further thank R. Benton for continuous support throughout this project, members of the Benton lab for constructive feedback and discussion, and S. Sprecher for sharing fly room space and comments on the manuscript. This work was supported by funding from FCT–Fundação para a Ciência e Tecnologia (PTDC/MED-NEU/4001/2021), the “la Caixa” Banking Foundation (project codes HR17-00539 and HR23-00516), the Champalimaud Foundation (supporting research at the Centre for the Unknown) (C.R.), a Deutsche Forschungsgemeinschaft Walter-Benjamin Fellowship (E.B.), a SNSF Ambizione (PZ00P3 185743) and Starting Grant (TMSGI3_211391/1), the Fondation Pierre Mercier pour la science, the Novartis Foundation for biomedical research, and the University of Fribourg (T.O.A.).

## Methods

### Data reporting

Preliminary experiments were used to assess variance and determine adequate sample sizes in advance of acquisition of the reported data. Several experiments were carried out repeatedly because they served as controls for different genetic manipulations. For calcium imaging, data were collected from multiple flies on multiple days in randomised order. Within datasets the same tastant dilutions were used for acquisition of the dataset. The experimenter was not blinded to the genotype of flies in physiological experiments.

### Drosophila husbandry

*Drosophila* stocks in the Auer lab were maintained on standard wheat flour/yeast/fruit juice medium under a 12 h light:12 h dark cycle at 22-25°C. For all *D. sechellia* strains, a few g of Formula 4-24® Instant *Drosophila* Medium, Blue (Carolina Biological Supply Company) soaked in noni juice (Raab Vitalfood) were added on top of the standard food. Flies used for volumetric imaging were reared on a yeast-based medium (containing per liter of water: 8 g agar (NZYTech), 80 g barley malt syrup (Próvida), 22 g sugar beet syrup (Grafschafter), 80 g cornflour (Próvida), 10 g soya flour (A. Centazi), 18 g instant yeast (Saf-instant, Lesaffre), 8 ml propionic acid (Argos), and 12 ml nipagin (Tegospet, Dutscher) (15% in 96% ethanol)). Food was supplemented with instant yeast granules on the surface (Saf-instant, Lesaffre) and stocks were kept at 25°C, 70% relative humidity in a 12 h light:12 h dark cycle. Wild-type, published and newly created transgenic lines used in this study are listed in **Sup. Table 2**.

### *Drosophila* microinjections

*D. sechellia* transgenesis was performed in-house as described^31^. For CRISPR-Cas9-mediated homologous recombination, a mix of a sgRNA-encoding construct (150 ng/ul) and donor vector (400 ng/ul) was injected into *Dsec nanos-Cas9*^31^. Site-directed integration into attP sites was achieved by co-injection of pBS130 (400 ng/ul) (encoding *PhiC31 integrase* under control of a heat shock promoter (Addgene #26290)^75^) and an *attB*-containing vector (400 ng/ul). All concentrations are given as final values in the injection mix.

### CRISPR/Cas9-mediated genome engineering and molecular cloning

#### sgRNA expression vectors

For expression of single sgRNAs, oligonucleotide pairs (**Sup. Table 3**) were annealed and cloned into *BbsI*-digested *pCFD3-dU6-3gRNA* (Addgene #49410)^76^. To express multiple sgRNAs from the same vector backbone, oligonucleotide pairs (**Sup. Table 4**) were used for PCR and inserted into *pCFD5* (Addgene #73914)^77^ via Gibson Assembly.

#### Donor vectors for homologous recombination

To target the *orco* locus of *D. simulans* a similar strategy as for the *orco^1^* allele in *D. sechellia* was applied^31^. Homology arms were amplified from *D. simulans* genomic DNA and inserted via restriction cloning into *pHD-3P3-DsRed*. To generate Gal4 reporter alleles, a 2A-Gal4 expression vector (GeneScript) was synthesized and either *D. sechellia*, *D. simulans* or *D. melanogaster* specific homology arms generated via gene synthesis (GeneScript) were introduced via seamless cloning flanking the expression cassette. Sequence details can be obtained from the corresponding author upon request.

#### UAS-expression vectors

cDNA vectors for *D. melanogaster Gr39a.a*, *Gr39a.b* and *D. sechellia Gr39a.a* were generated via gene synthesis (GeneScript). cDNA was amplified via PCR and inserted into *pUAST-attB*^78^ via restriction cloning resulting in *pUAS-DmelGr39a.a-attB, pUAS-DmelGr39a.b-attB*, and *pUAS-DsecGr39a.a-attB*.

#### Site directed mutagenesis

A three amino acid deletion was introduced into *D. melanogaster pUAS-DmelGr39a.a-attB* resulting in *pUAS-DmelGr39a.a^Δ3^-attB* via PCR mediated site directed mutagenesis (sequences of oligonucleotides are listed in **Sup. Table 4**).

### Immunohistochemistry

Immunofluorescence on adult brains, proboscis, and legs was performed as previously described^79^ using mouse monoclonal antibody nc82 1:10 (Developmental Studies Hybridoma Bank), anti-Ir25a (guinea pig, gift from R. Benton), and rabbit anti-GFP 1:500 (Invitrogen). Alexa488-, Cy3- and Cy5- conjugated goat anti-guinea pig, goat anti-mouse, and goat anti-rabbit IgG secondary antibodies (Molecular Probes, Jackson Immunoresearch) were used at 1:500.

### Image acquisition and processing

Confocal images of brains, legs and proboscis were acquired on an inverted confocal microscope (Zeiss LSM 710) equipped with an oil immersion 40Å∼ objective (Plan Neofluar 40Å∼ oil immersion DIC objective; 1.3 NA) and images were processed in Fiji^80^. *D. sechellia* brains were registered to a *D. sechellia* reference brain^31^ using the Fiji CMTK plugin (https://github.com/jefferis/fiji-cmtk-gui), as previously described^81^.

### Behavioural assays

#### 4-choice arena

A group of 20 male and 50 female flies was anaesthetized 5 days after eclosion on ice and transferred to an arena (20x20x5 cm) containing four different fruits (pieces of apple, banana, grapes, and noni) placed on a small petri dish (6 cm diameter). The arena was covered with a mesh, placed under a camera and snapshots were take every 10 minutes for 24 hours. Apples, banana and grape were purchased freshly from a local supermarket; noni at the ripe stage was harvested from the greenhouse at the University of Lausanne. Assays were run at red light under temperature (25°C) and humidity (60%) controlled conditions starting at 11am for each replicate. The position of fruits was randomized for each experiment. At every timepoint, the number of flies per quadrant was counted; data is represented as percentage of flies per quadrant.

#### 72-well plate feeding assay

The 72-well plate feeding assay was adapted from previous studies^11,79^. Briefly, agarose (1%) was disolved in noni or grape juice, heated in a microwave and mixed with food colorants (Food Red No. 106; 500 μg/ml; Food Blue No. 1, 125 μg/ml; Tokyo Chemical Industry Co., Tokyo, Japan). 10 μl of each solution were pipetted into alternating wells of 72-well microtiter plates (Sigma M5812). Groups of 80 male or female 3-7 day old mated flies were wet-starved for 22h at 22°C, anaesthetized on ice for 1 minute and transferred into the plates where they were allowed to feed for 1.5 hours in darkness at 25°C, 60% relative humidity. Flies’ feeding was terminated by plate flash freezing and food ingestion was scored by two independent operators. Juice-to-colourant association was swapped at every replicate. The percentage of feeding was calculated by scoring flies without colourant in the abdomen and comparison to flies with food consumption. The preference index was calculated with the following formula: (number of flies feeding on A - number of flies feeding on B)/total number of flies feeding.

#### Petri dish feeding assay

The petri dish feeding assay was adapted from previous reports^32,82^. Briefly, groups of 20 male or female 3-7 day old mated flies were wet-starved for 22 h at 22°C, anaesthetized on ice for 1 minute and transferred into small Petri dishes (60x15 mm, Sarstedt 82.1194.500) where food substrates in 1.3% moderate EEO-agarose (SeaKem ME agarose, CatNo 50010) labelled with blue or red colourants (as above) were pipetted as 18 (9x2) semisolid drops (10 μl) on the dish surface. A template was used for reproducible and equally spaced spot distribution on the petri dish surface. For experiments with antennectomized flies, surgery was performed on freshly eclosed flies followed by 3-5 days of recovery before starvation. The preference index was calculated as for the 72-well plate feeding assay.

#### FlyPAD

The flyPAD (fly Proboscis and Activity Detector) rig was purchased from EasyBehaviour and used in a two-choice configuration where food substrates were pipetted into the sensor wells as 1.3% moderate EEO-agarose solutions (as for the petri dish assay). Starved flies (22h wet starvation at 22°C) from different genetic backgrounds were loaded in the arena using a mouth aspirator in random order and allowed to feed under dim red light for 1 hour at 25°C, 60% relative humidity. The flyPAD capacitance signal was analysed as previously reported^33^ and multiple feeding parameters were extracted. The preference index was calculated as (number of sips on food A - number of sips on food B)/total number of sips in 1 hour. Principal component analysis was performed including all flyPAD extracted parameters using the *prcomp()* function in R.

#### Proboscis extension response (PER)

PER was assessed on starved female flies (22 h wet starvation at 22°C) mounted inside a pipette tip so that only the head was exposed for labellar stimulation. After mounting, flies were allowed to recover in a humidified chamber for 1 hour and water was provided until satiation before testing. Sucrose (200 mM) was applied to the labellum with a blunted needle syringe containing the liquid tastant solution and stimulation was repeated for three times with a one-minute interval between stimulations. PER was scored when a fly showed full proboscis extension, and the experimenter was blind to the genotype of tested flies.

#### Temporal consumption assay

The temporal consumption/proboscis extension time of flies was quantified as previously described^64^ with slight modifications: 3-5 day old females flies were wet-starved at 22°C for 22 h, anesthetized using CO_2_ and fixed to a glass slide with glue (Pritt). Flies recovered for 1-2 hrs in a humidified box and were given water ad libitum prior to testing with other tastants. In testing, flies were presented with one tastant (200 mM sucrose, grape juice or noni juice) 10 times and consumption/proboscis extension time was manually recorded for each fly and summed up. Stimuli order was randomized. If flies did not respond to a tastant they were tested for response to any of the other two and only included if they responded to at least one of the stimuli. All experiments were done in light, at constant temperature (25°C) and humidity, and the experimenter was blind to the genotype of tested flies.

### Calcium imaging

#### Widefield imaging

SEZ imaging was performed as previously described^50^. For all experiments, 3-7 old non-starved female flies raised in groups at 22°C were used. These were anaesthesized on ice, mounted into home-made imaging blocks with extended proboscis and fixed with glue (UV glue, Tetric EvoFlow, A1, Ivoclar Vivadent). The proboscis was sealed from the head capsule to prevent buffer leakage. The head was surgically opened frontally by removing the antennae and the SEZ was exposed to the microscope objective in AHL buffer (Adult Haemolymph-Like saline buffer: 108 mM NaCl, 5 mM KCl, 2 mM CaCl2, 8.2 mM MgCl2, 4 mM NaHCO3, 1 mM NaH2 PO4, 5 mM trehalose, 10 mM sucrose, 5 mM HEPES, pH 7.5) after removal of the oesophagus. A CCD camera (CoolSNAP-HQ2 Digital CameraSystem) mounted on a fluorescence microscope (upright fixed stage Carl Zeiss Axio Examiner D1) equipped with a 40x water-immersion objective (W Plan-Apochromat 40Å∼ /1,0 VIS-IR DIC) was used for GFP signal acquisition. The excitation light (470 nm) was produced with an LED light (Cool LED pE-100, VisiChrome). Tastants were delivered manually using a micromanipulator (Narishige) equipped with a blunted needle syringe containing the liquid tastant solution. Stimulation was initiated 10 seconds after the LED was turned on and maintained for 10 seconds, after which contact was terminated while recording continued for an additional 10 seconds. The order of different tastant application was partly randomized: water was always applied first followed by other tastants. Water was applied also in between stimulation to wash away residues from previous applications on the labellum, though these water applications were not quantified. Juice solutions were used fresh from frozen aliquots. The other tastants were re-used for a maximum of 3 months. Time course images (16-bit) were acquired using Metafluor software (Visitron) at a rate of 3 Hz with a 200 ms exposure time per frame. The signal time course was quantified using Fiji^80^ by extracting the maximum pixel value over time from manually selected ROIs. ROIs were drawn around SEZ projections and background areas and the average background intensity was subtracted from SEZ projections intensity. Basal fluorescence (F) was determined by averaging the intensity of 10 timepoints prior to the onset of the stimulation, while peak fluorescence was identified as the maximum intensity within the 10 timepoints following the onset. The relative changes in fluorescence (ΔF/F) were then calculated. To account for species-specific differences in GCaMP levels and to enable inter-species comparisons, we normalised the relative changes in fluorescence for each tastant application by the average response within the same preparation.

#### Volumetric imaging

Pan-neuronal volumetric imaging in the SEZ was performed as previously described^18^. 1-15d old female flies were cold-anaesthesized on ice and fixed to a custom-made imaging block using fast UV curing glue (Bondic). The proboscis was fixed in an extended position and the front legs were removed to prevent the flies from touching the food stimulus. The anterior part of the head was isolated from the rest of the body through a hole in a weigh ship and covered with *Drosophila* AHL (103 mM NaCl, 3 mM KCl, 5 mM TES, 10 mM trehalose dihydrate, 10 mM glucose, 2 mM sucrose, 26 mM NaHCO3, 2 mM CaCl2 dihydrate, 4 mM MgCl 2 hexahydrate, 1 mM NaH2PO4, pH 7.3). A window was cut between the eyes and the ocelli, thereby removing the antennae, the oesophagus was transected to allow for unoccluded visual access to the SEZ.

A resonant-scanning two-photon microscope (Scientifica) equipped with a 20×NA 1.0 water immersion objective (Olympus) and a piezo-electric z-focus drive, was used to allow for fast volumetric scans. SEZ volumes in the three different species were acquired for 60 s at a 1 Hz volume rate covering 512 × 256 × 60 voxels at voxel dimensions of ∼0.5 × 0.5 × 3.6 μm using SciScan (Scientifica). A Chameleon Ultra II Ti:Sapphire laser (Coherent) was used to excite GCaMP6s at 920 nm. During imaging, the brain was constantly perfused with AHL bubbled with carbogen (95% O_2_, 5% CO_2_). Manual gustatory stimulations of the proboscis were performed using glass capillaries filled with taste solution, mounted on a micromanipulator (Sensapex).

Species-specific brain templates were generated using the Advanced Normalization Toolkit (ANTs) script antsMultivariateTemplateConstruction2.sh. We corrected for movement artifacts and shifts between recordings within an animal using ANTs. Recordings were then pre-aligned to the respective species template by using manual landmarks that were defined in ITK-SNAP^83^. Finally, ANTs was used for automated fine registration of each recording to the respective template. We created average brain volumes for each stimulus-species combination and performed a maximum response projection. These volumes were used to define 3D ROIs in ITK-SNAP, which were then used for time course extraction from each recording. The ROIs included the anterior motor region, the sensory projections of bristle sensory neurons (PMS2/3 and PMS4), and two broad interneuron regions. Data quantification was performed as explained above.

### AlphaFold structural prediction

*D. sechellia* Gr39a.a peptide sequence was translated from aligned nucleotide sequences with *D. melanogaster*, protein structure predicted using ColabFold^84^ and plotted using PyMOL (Version 1.8 Schrödinger, LLC).

### Fly Cell Atlas analysis

Single nuclear transcriptomic data were obtained from the Fly Cell Atlas^85^ datasets (s_fca_biohub_proboscis_and_maxillary_palps_10x, s_fca_biohub_antenna_10x) and transcript co-expression was determined without expression threshold.

### Statistics and reproducibility

Data were analysed and plotted using Excel, R (v4.4.1; R Core Team 2024) and the tidyverse suite^86^. All statistical details are listed in the figures and figure legends and exact p-values are listed in **Sup. Table 5**.

### Data, code and biological material availability

All relevant data supporting the findings of this study, code used for analysis and all unique biological materials generated in this study (e.g., mutants, plasmids and transgenic fly strains) are available from the corresponding author upon request.

## Figure Legends

**Supplementary Figure 1: Feeding preference is not sexually dimorphic**

**A)** Distribution of flies per quadrant in the 4-choice assay from 10 min to 24 h after assay start, seperated by species.

**B)** Feeding preference for female (left) and male (right) flies in the 72-well, petri dish and flyPAD assay. *D.sechellia* = DSSC 14021-0248.07 and 14021-0248.28. *D. simulans* = DSSC 14021-0251.196 and DSSC 14021-0251.004. *D. melanogaster* = *Dmel CantonS and Oregon R* (here and in other panels). Data is re-plotted from Fig. 1. Here and in other panels, sample sizes are indicated in the figure. Kruskal-Wallis H test (72-well, females: H_(5)_=60.22, P=1.095e^-11^; 72-well, males: H_(5)_=29.109, P=2.207e^-05^; petri dish, females: H_(5)_=35.773, P=1.055e^-06^; petri dish, males: H_(5)_=33.573, P=2.895e^-06^; flyPAD, females: H_(5)_=217.2, P=2.2e^-16^; flyPAD, males: H_(5)_=31.49, P=7.496e^-06^). Pairwise comparisons were conducted using Dunn’s test with Bonferroni correction, for these and all other comparisons the results are shown in the figure, where groups sharing the same letter are not significantly different (P < 0.025).

**C)** Percentage of female (left) and male (right) flies feeding on either noni juice, grape juice or both in the 72-well (two panels to the left) and petri dish assay (two panels to the right).

**D)** Number of sips within one hour after assay start on noni or grape juice in the flyPAD assay. The same data was used to calculate the preference index shown in Fig. 1B.

**E)** Feeding preference of female flies lacking eye pigmentation for noni or grape juice in the flyPAD assay. *D. melanogaster* = *Dmel w^1118^, D. simulans* = DSSC 14021-0251.195, *D.sechellia* = DSSC 14021-0248.30). Kruskal-Wallis H test (H_(2)_=77.005, P=2.2e^-16^; Dunn’s test results are shown in the figure).

**Supplementary Figure 2: *DsimOrco^1-/-^* mutant generation and flyPAD analysis**

**A)** Left: Schematic of the *Dsim orco* locus. The *3P3-DsRed* marker was inserted into the first exon to match the *DsecOrco^1^* allele^31^. In red: location of the epitope recognized by an Orco-specific antibody. Note that this allele represents a strong hypomorph but does not completely disrupt Orco function^31^. Right: Anti-Orco immunofluorescence in wild-type (DSSC 14021-0251.04, left) and *DsimOrco*^1*-/-*^ mutant antenna (right). Orco expression is dramatically reduced but not completely absent in *DsimOrco*^1^. Scale bar = 20 µm.

**B)** Left: Preference index of antennal removed female (left) or male (right) flies for noni vs. grape juice feeding in the petri-dish assay. Data is replotted from Fig. 1. Kruskal-Wallis H test (females: H_(2)_=13.62, P=0.001102; males: H_(2)_=5.4873, P=0.06434). Pairwise comparisons were conducted using Dunn’s test with Bonferroni correction, for these and all other comparisons the results are shown in the figure, where groups sharing the same letter are not significantly different (P < 0.025). Right: Percentage of feeding on either noni juice, grape juice or both separated by sex for flies shown to the left.

**C)** Schematic of the feeding parameters analysed with the flyPAD. Adapted from Ref.^33^.

**D)** Loadings of principal component 1 (left) and 2 (right) from Fig. 1E.

**E)** Sugar content of apple, banana, grape, grape juice, noni juice and noni fruit. Data was taken from the U.S. Department of Agriculture, Agricultural Research Service FoodData Central Database (https://fdc.nal.usda.gov/, identifiers: banana (9040), apple (9050), grapes (100280)) or the Malaysian Food Composition Database (https://myfcd.moh.gov.my/, identifier: *Morinda citrifolia* (105056)). Percentages are calculated based on g/100 ml for juices and g/100 g for fruits.

**F)** Percentage of feeding on either noni, sucrose or both. Data is replotted from Fig. 1F.

**G)** Number of sips per substrate as measured in the flyPAD assay after silencing of defined gustatory neuron populations. The same data was used to calculate the preference index shown in Fig. 1G.

**Supplementary Figure 3: Sweet sensory neuron population analysis**

**A)** Schematic illustrating the CRISPR/Cas9-mediated knock-in strategy to generate reporter *Gal4*-alleles at receptor loci.

**B)** Expression of Ir25a and Gr64f in individual neurons of the fly proboscis and maxillary palp (left) or fly antenna (right). Data from the FlyCellAtlas^85^.

**C)** Immunofluorescence in the labellum of *DsecGr64f^Gal4^>UAS-GCaMP6s* transgenic flies with an anti-GFP (detecting GCaMP6s expression) and anti-Ir25a antibody. Gr64f expressing cells represent a subpopulation of Ir25a expressing neurons. Scale bar = 20 µm.

**D)** Number of Gr64f^+^ neurons in the labellum and legs of male *D. melanogaster*, *D. simulans* and *D. sechellia* based on transgenic labelling. Here and in other panels, sample size is indicated in the figure. For these and all other cell counts bar plots, the bar represents the mean, the error bar the standard error; individual data points are overlaid. Kruskal-Wallis H test (labellum: H_(2)_=14.05, P=0.0008894; foreleg: H_(2)_=17.631, P=0.0001484; midleg: H_(2)_=6.0282, P=0.04909; hindleg: H_(2)_=1.3862, P=0.5). Dunn’s test results (after Bonferroni correction) are shown in the figure.

**E)** Non-normalised calcium imaging data in taste pegs (same as Fig. 2E). Wilcoxon rank-sum. No significant differences were detected.

**F)** Non-normalised calcium imaging data in taste bristles (same as Fig. 2F). Wilcoxon rank-sum test. (*P < 0.05). Significant differences to *D. melanogaster* responses are shown in the figure.

**Supplementary Figure 4: Bitter sensory neuron characterization**

**A)** Number of Gr66a^+^ neurons in the labellum and legs of male *D. melanogaster*, *D. simulans* and *D. sechellia* based on transgenic labelling. Here and in other panels, sample size is indicated in the figure. For these and all other cell counts bar plots, the bar represents the mean, the error bar the standard error; individual data points are overlaid. Kruskal-Wallis H test (labellum: H_(2)_=14.05, P=0.0008894; foreleg: H_(2)_=17.631, P=0.0001484; midleg: H_(2)_=6.0282, P=0.04909; hindleg: H_(2)_=1.3862, P=0.5). Dunn’s test results (after Bonferroni correction) are shown in the figure.

**B)** Non-normalised calcium imaging data in taste bristles (same as Fig. 3C).

**C)** Quantification of tastant evoked calcium responses in Gr66a^+^ bristles neurons of the fly labellum in the SEZ of *D. melanogaster*, *D. simulans*, and *D. sechellia* towards water, sucrose, caffeine and denatonium. This dataset is independent from data shown in Fig. 3. Wilcoxon rank-sum test. (***P < 0.001). Significant differences to *D. melanogaster* responses are shown in the figure.

**Supplementary Figure 5: Physiology of *DmelGr39a.b* rescue and volumetric imaging data in *D. simulans***

**A)** Quantification of tastant evoked calcium responses in Gr66a^+^ bristles neuron projections in the SEZ of *D. sechellia* expressing *DmelGr39a.b*. Wilcoxon rank-sum test. No significant differences to wild-type are detected.

**B)** Preference index for denatonium (10 mM) in 50 mM sucrose vs. 5 mM sucrose feeding in the petri dish assay for all three species. The variance of the data = 0.

**C)** Temporal fluorescence changes in sweet/other axonal innervations in *D. melanogaster* and *D. sechellia* upon water and denatonium benzoate (10mM) presentation. The black bars indicate the interval of tastant application.

**D)** Quantification of tastant evoked neuronal activity (maximum responses) in the segmented areas shown in Fig. 3B in *D. simulans.* For visualization purposes, the following outliers are excluded from the display but included in the statistical analysis (*D. simulans*/interneurons-1/noni juice ΔF/F= −139%; *D. simulans*/interneurons-2/water ΔF/F= 401%; *D. simulans*/bitter/grape juice ΔF/F= - 267%). Wilcoxon rank-sum test. (*P < 0.05). Significant differences to *D. melanogaster* responses are shown in the figure.

