## Supplementary Table 2,3,4 for "Evolution of taste processing shifts dietary preference"

**Supplementary Table 1:** Number of Gr64fa- and Gr66a-expressing neurons in *D. melanogaster*, *D. simulans* and *D. sechellia* taste organs.

**Supplementary Table 2.** Wild-type and transgenic lines used in this study.

| **Stock name** | **Donor plasmid** | **Parental strain** | **Species** | **Method/Reference** | **Figure** |
|---|---|---|---|---|---|
| *Dsec.07* | | *Drosophila* Species Stock Center [DSSC] 14021-0248.07 | *D. sechellia* | | Fig. 1, 2, 4, 5; Sup. Fig. 1, 2, 5 |
| *Dsec.30* | | DSSC 14021-0248.30 | *D. sechellia* | | Fig. 1; Sup. Fig. 1 |
| *Dsec.28* | | DSSC 14021-0248.28 | *D. sechellia* | | Fig. 1; Sup. Fig. 1 |
| *Dmel CS* | | *D. melanogaster Canton S* | *D. melanogaster* | | Fig. 1, 2, 4, 5; Sup. Fig. 1, 2, 5 |
| *Dmel OR* | | *D. melanogaster Oregon R* | *D. melanogaster* | | Fig. 1; Sup. Fig. 1 |
| *Dmel* $w^{1118}$ | | *D. melanogaster w1118* | *D. melanogaster* | | Fig. 1; Sup. Fig. 1, 2 |
| *Dsim.04* | | DSSC 14021-0251.004 | *D. simulans* | | Fig. 1, 2, 4, 5; Sup. Fig. 1, 2, 5 |
| *Dsim.196* | | DSSC 14021-0251.196 | *D. simulans* | | Fig. 1; Sup. Fig. 1 |
| *Dsim.195* | | DSSC 14021-0251.195 | *D. simulans* | | Fig. 1; Sup. Fig. 1 |
| $DsecIr8a^{-/-}/Orco^{1-/-}$ | | | *D. sechellia* | [31] | Fig. 1; Sup. Fig. 1 |
| $DsecOrco^{1}$ | | | *D. sechellia* | [31] | Fig. 1; Sup. Fig. 2 |
| $DsimOrco^{1}$ | *pHD-DsimOrco-3xP3-DsRed* | DSSC 14021-0251.195 | *D. simulans* | CRISPR knock-in, this study | Fig. 1; Sup. Fig. 2 |
| $DmelOrco^{-/-}$ | | BDSC_23130 | *D. melanogaster* | | Fig. 1; Sup. Fig. 2 |
| *Dmel* $Ir8a^{-/-}/Orco^{-/-}/Ir25a^{-/-}/Gr63a^{-/-}$ | | | *D. melanogaster* | [87] | Fig. 1; |
| *Dmel Gr64f-Gal4* | | Bloomington Drosophila Stock Center [BDSC] 57669 | *D. melanogaster* | | Fig. 1, 2; Sup. Fig. 2, 3 |
| *Dmel Gr66a-Gal4* | | | *D. melanogaster* | [88] | Fig. 1, 3; Sup. Fig. 2 |
| *Dmel Ir25a-Gal4* | | BDSC_41728 | *D. melanogaster* | | Fig. 1, 2; Sup. |

| | | | | | |
|---|---|---|---|---|---|
| | | | | | Fig. 2, 3 |
| *Dmel ppk28-Gal4* | | BDSC_93020 | *D. melanogaster* | | Fig. 1; Sup. Fig. 2 |
| *Dmel ppk23-Gal4* | | BDSC_93026 | *D. melanogaster* | | Fig. 1; Sup. Fig. 2 |
| *Dmel Ir94e-Gal4* | | BDSC_81246 | *D. melanogaster* | | Fig. 1; Sup. Fig. 2 |
| *Dmel UAS-Kir2.1* | | BDSC_6595 | *D. melanogaster* | | Fig. 1; Sup. Fig. 2 |
| *DsecGr66a$^{Gal4}$* | *pHD-DsecGr66a-T2A-Gal4-3xP3-DsRed* | *Dsec*nanos-Cas9[31] | *D. sechellia* | CRISPR knock-in, this study | Fig. 3, 4; Sup. Fig. 4, 5 |
| *DsecGr64f$^{Gal4}$* | *pHD-DsecGr64f-T2A-Gal4-3xP3-DsRed* | *Dsec*nanos-Cas9[31] | *D. sechellia* | CRISPR knock-in, this study | Fig. 2; Sup. Fig. 3 |
| *DsecIr25a$^{Gal4}$* | *pHD-DsecIr25a-T2A-Gal4-3xP3-DsRed* | *Dsec*nanos-Cas9[31] | *D. sechellia* | CRISPR knock-in, this study | Fig. 2; Sup. Fig. 3 |
| *Dsec unc84 ::GFP* | *pMUH_unc84_2xGFP (Addgene #46023)* | *Dsec-pBac-attPA26*[89] | *D. sechellia* | attB/P integration, this study | Fig. 2, 3; Sup. Fig. 3, 4 |
| *DmelGr66a$^{Gal4}$* | *pHD-DmelGr66a-T2A-Gal4-3xP3-DsRed* | | *D. melanogaster* | CRISPR knock-in, this study | Fig. 3; Sup. Fig. 4 |
| *DmelGr64f$^{Gal4}$* | *pHD-DmelGr64f-T2A-Gal4-3xP3-DsRed* | | *D. melanogaster* | CRISPR knock-in, this study | Fig. 2; Sup. Fig. 3 |
| *Dmel unc84 ::GFP* | | | *D. melanogaster* | [35] | Fig. 2, 3; Sup. Fig. 3, 4 |
| *DsimGr66a$^{Gal4}$* | *pHD-DsimGr66a-T2A-Gal4-3xP3-DsRed* | | *D. simulans* | CRISPR knock-in, this study | Fig. 3; Sup. Fig. 4 |
| *DsimGr64f$^{Gal4}$* | *pHD-DsimGr64f-T2A-Gal4-3xP3-DsRed* | | *D. simulans* | CRISPR knock-in, this study | Fig. 2; Sup. Fig. 3 |
| *Dsim unc84 ::GFP* | *pMUH_unc84_2xGFP (Addgene #46023)* | *Dsim#2176*[90] | *D. simulans* | attB/P integration, this study | Fig. 2, 3; Sup. Fig. 3, 4 |
| *DsecUASGCaMP6s* | | | *D. sechellia* | [89] | Fig. 2, 3, 4, 5; Sup. Fig. 3, 4, 5 |
| *DmelGr39a$^{-/-}$* | | | *D. melanogaster* | [25] | Fig. 4 |
| *Dsec-UAS-DmelGr39a.a* | *attB-DmelGr39a.a* | *Dsec-pBac-attPA26*[89] | *D. sechellia* | attB/P integration, this study | Fig. 4 |
| *Dsec-UAS-DmelGr39a.b* | *attB-DmelGr39a.b* | *Dsec-pBac-attPA26*[89] | *D. sechellia* | attB/P integration, this study | Fig. 4; Sup. Fig. 5 |
| *Dmel Gr39a.a-Gal4* | | BDSC_57631 | *D. melanogaster* | [91] | Fig. 4 |
| *Dmel-UAS-DmelGr39a.a* | *attB-DmelGr39a.a* | BDSC_24749 | *D. melanogaster* | attB/P integration, this study | Fig. 4 |
| *Dmel-UAS-DsecGr39a.a* | *attB-DsecGr39a.a* | BDSC_24749 | *D. melanogaster* | attB/P integration, this study | Fig. 4 |
| *Dmel-UAS-DmelGr39a.a $^{\Delta 3}$* | *attB-DmelGr39a.a $^{\Delta 3}$* | BDSC_24749 | *D. melanogaster* | attB/P integration, this study | Fig. 4 |
| *DsecNsyb-Gal4* | | | *D. sechellia* | [31] | Fig. 5; Sup. Fig. 5 |
| *DsimNsyb-Gal4* | | | *D. simulans* | [63] | Sup. |

| | | | | | |
|---|---|---|---|---|---|
| | | | | | Fig. 5 |
| *DmelNsyb-Gal4* | ;nSyb-Gal4 3-1 (2);; | | *D. melanogaster* | Gift from Julie Simpson | Fig. 5; Sup. Fig. 5 |
| *Dsim UAS-GCaMP6s* | | | *D. simulans* | [90] | Sup. Fig. 4, 5 |
| *Dmel UAS-GCaMP6s* | | BDSC_42746 | *D. melanogaster* | | Fig. 2, 3, 4, 5; Sup. Fig. 3, 4, 5 |

**Supplementary Table 3.** Oligonucleotides used to generate sgRNA expression vectors.

| Target | Name | Sequence (5'-3') | Resulting sgRNAs |
|---|---|---|---|
| *DsimOrco* | *PCR1fwd* | GCGGCCCGGGTTCGATTCCCGGCCGATGCAGCCGAGCAAGTACACGGGCCGTTTTAGAGCTAGAAATAGCAAG | sgRNA1: GCCGAGCAAGTACACGGGCC |
| | *PCR1rev* | CTCGACCCAGTACTTGATGGTGCACCAGCCGGGAATCGAACCC | |
| | *PCR2fwd* | CCATCAAGTACTGGGTCGAGGTTTTAGAGCTAGAAATAGCAAG | sgRNA2: CCATCAAGTACTGGGTCGAG |
| | *PCR2rev* | CCCCCGTTACCGCCGAAGGATGCACCAGCCGGGAATCGAACCC | |
| | *PCR3fwd* | TCCTTCGGCGGTAACGGGGGGTTTTAGAGCTAGAAATAGCAAG | sgRNA3: TCCTTCGGCGGTAACGGGGG |
| | *PCR3rev* | ATTTTAACTTGCTATTTCTAGCTCTAAAACGCCACCGACATGGACATGTCTGCACCAGCCGGGAATCGAACCC | sgRNA4: GACATGTCCATGTCGGTGGC |
| *DmelGr66a* | *PCR1fwd* | GCGGCCCGGGTTCGATTCCCGGCCGATGCAGATAGCTGGAATTCTGCCACGTTTTAGAGCTAGAAATAGCAAG | sgRNA1: GATAGCTGGAATTCTGCCAC |
| | *PCR1rev* | ATTTTAACTTGCTATTTCTAGCTCTAAAACTTACGGGATTTCTCCAGCAGTGCACCAGCCGGGAATCGAACCC | sgRNA2: CTGCTGGAGAAATCCCGTAA |
| *DsimGr66a* | *PCR1fwd* | GCGGCCCGGGTTCGATTCCCGGCCGATGCACTGCTGGAGAAATCCCGTAAGTTTTAGAGCTAGAAATAGCAAG | sgRNA1: CTGCTGGAGAAATCCCGTAA |
| | *PCR1rev* | ATTTTAACTTGCTATTTCTAGCTCTAAAACTTACGGGATCTCTCCAGCAGTGCACCAGCCGGGAATCGAACCC | sgRNA2: CTGCTGGAGAGATCCCGTAA |
| *DsecGr66a* | *PCR1fwd* | GCGGCCCGGGTTCGATTCCCGGCCGATGCAGATAGCTGGAATTCTGCCGCGTTTTAGAGCTAGAAATAGCAAG | sgRNA1: GATAGCTGGAATTCTGCCGC |
| | *PCR1rev* | ATTTTAACTTGCTATTTCTAGCTCTAAAACTTACGGGATCTCTCCAGCAGTGCACCAGCCGGGAATCGAACCC | sgRNA2: CTGCTGGAGAGATCCCGTAA |
| *DmelGr64f* | *PCR1fwd* | GCGGCCCGGGTTCGATTCCCGGCCGATGCAAAGATTCTTCCGAAGCTGGAGTTTTAGAGCTAGAAATAGCAAG | sgRNA1: AAGATTCTTCCGAAGCTGGA |
| | *PCR1rev* | ATTTTAACTTGCTATTTCTAGCTCTAAAACGTGATGTCCGGGTTACCCTTTGCACCAGCCGGGAATCGAACCC | sgRNA2: AAGGGTAACCCGGACATCAC |
| *DsimGr64f* | *PCR1fwd* | GCGGCCCGGGTTCGATTCCCGGCCGATGCAAAGATCCTTCCCAAGCTGGAGTTTTAGAGCTAGAAATAGCAAG | sgRNA1: AAGATCCTTCCCAAGCTGGA |
| | *PCR1rev* | ATTTTAACTTGCTATTTCTAGCTCTAAAACGTGATGTCCGGGTTACCCTTTGCACCAGCCGGGAATCGAACCC | sgRNA2: AAGGGTAACCCGGACATCAC |
| *DsecGr64f* | *PCR1fwd* | GCGGCCCGGGTTCGATTCCCGGCCGATGCACGCAACTTGCCCTCCAGCTTGTTTTAGAGCTAGAAATAGCAAG | sgRNA1: CGCAACTTGCCCTCCAGCTT |
| | *PCR1rev* | GAGATGTCCGAGTTACCCTTTGCACCAGCCGGGAATCGAACCC | sgRNA2: AAGGGTAACTCGGACATCTC |
| | *PCR2fwd* | AAGGGTAACTCGGACATCTCGTTTTAGAGCTAGAAATAGCAAG | sgRNA3: AAGATCCTTCCAAAGCTGGA |
| | *PCR2rev* | ATTTTAACTTGCTATTTCTAGCTCTAAAACTCCAGCTTTGGAAGGATCTTTGCACCAGCCGGGAATCGAACCC | |

| | | | |
|---|---|---|---|
| *DsecIr25a* | *PCR1fwd* | GCGGCCCGGGTTCGATTCCCGGCCGATGCAACTATACTTCGAATGGCCTCGTTTTAGAGCTAGAAATAGCAAG | sgRNA1: ACTATACTTCGAATGGCCTC |
| | *PCR1rev* | ATTTTAACTTGCTATTTCTAGCTCTAAAACGGGCCTGGGTGAAGCTCTTCTGCACCAGCCGGGAATCGAACCC | sgRNA2: GAAGAGCTTCACCCAGGCCC |

**Supplementary Table 4.** Oligonucleotides used for plasmid cloning.

| **Amplicon** | **Forward primer (5'-3')** | **Reverse primer (5'-3')** |
|---|---|---|
| *DmelGr39a.a* | GCCGCCCGACGCTTCGGCAAGCTCACTAAGGCGCCTGAGAAGAGCGGCGAACGGG | CCCGTTCGCCGCTCTTCTCAGGCGCCTTAGTGAGCTTGCCGAAGCGTCGGGCGGC |
| *DmelGr39a.b* | GATCGGATCCTCGACCATGGGCACAAGAAATCGAAAGCTTCTG | GATCCTCGAGTCAAAATTTATTAATAGACTTTGTGGA |
| *DmelGr39a.a*$^{\Delta 3}$ | CAACCAACAGCTTTGTCAGGCCTCTGGTCTTCGAGGCAGGCAGCAGATCC | CTGCCTCGAAGACCAGAGGCCTGACAAAGCTGTTGGTTGTAGTGAATCAATCCCGCGG |
| *DsecGr39a.a* | GATCGAATTCATGTCAAAAGTCTGCCGGG | GATCCTCGAGTCAAAATTTATTAATAGACTTTGTGG |

**Supplementary Table 5.** *P*-values of statistical analysis.
