## Supplementary figures and images for "Evolution of taste processing shifts dietary preference"

### Supplementary Figures

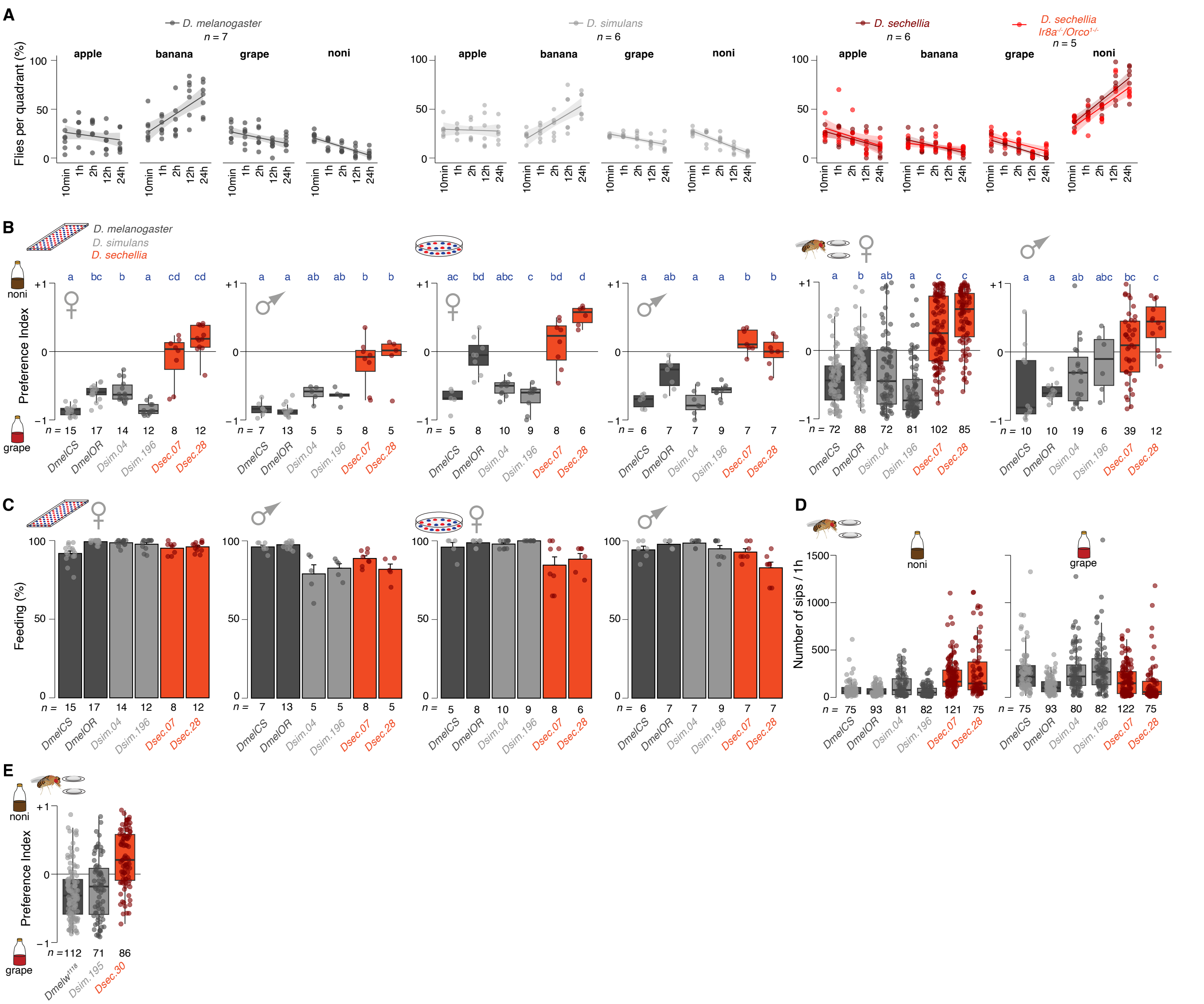

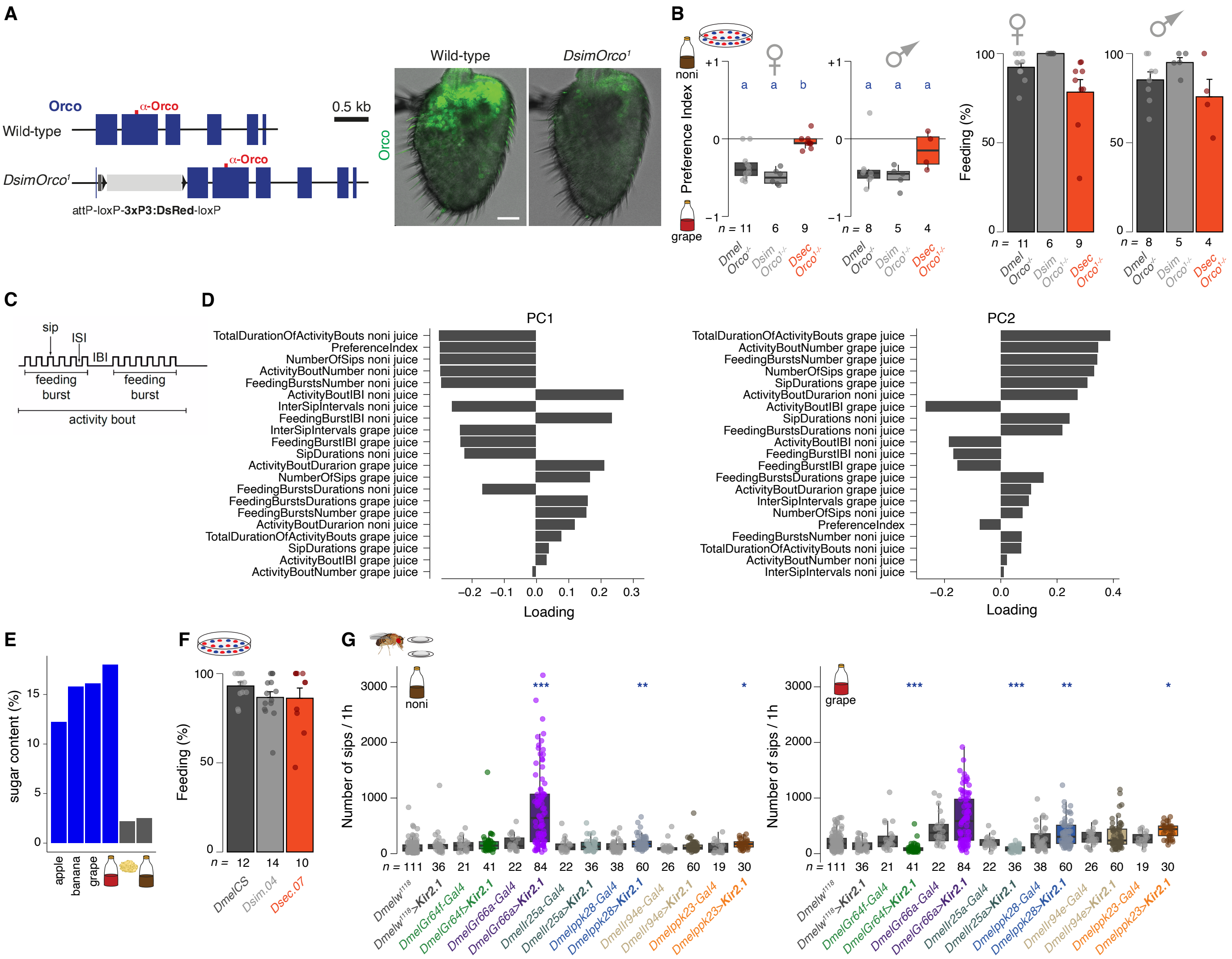

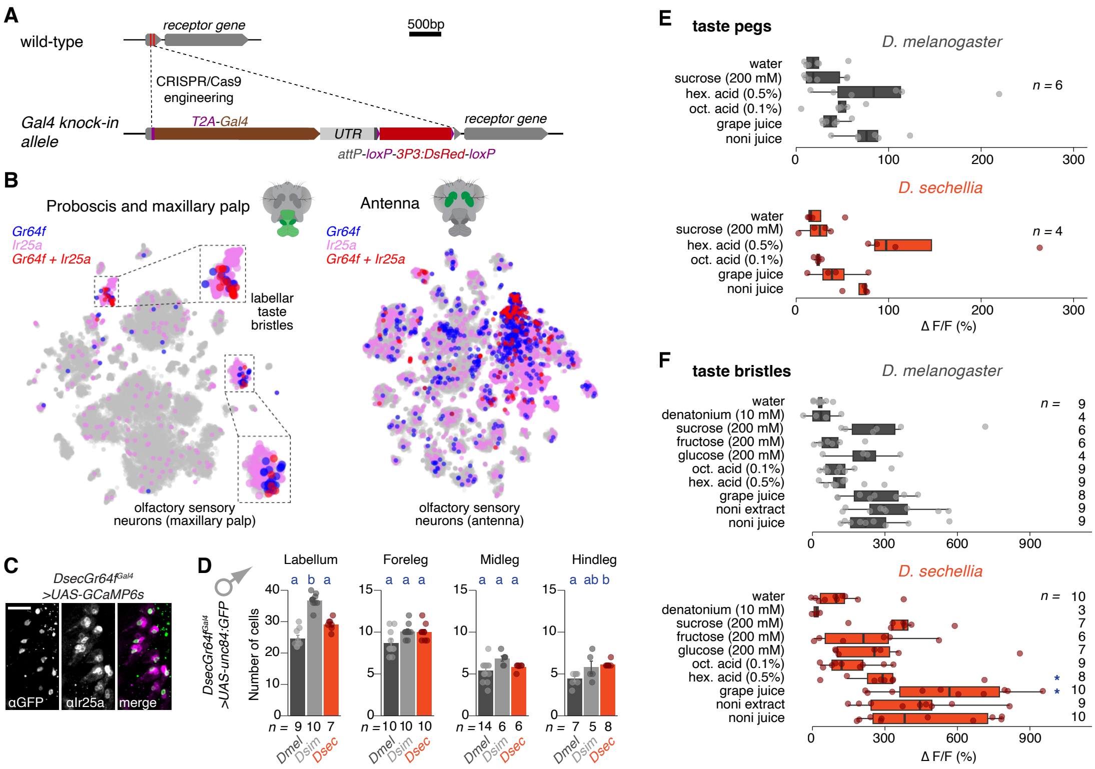

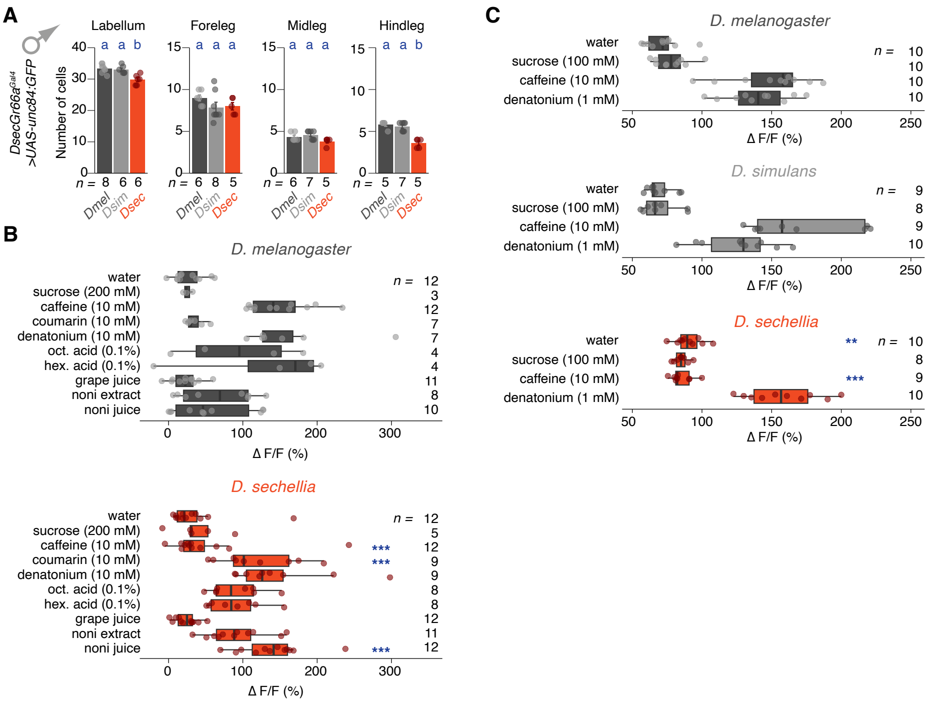

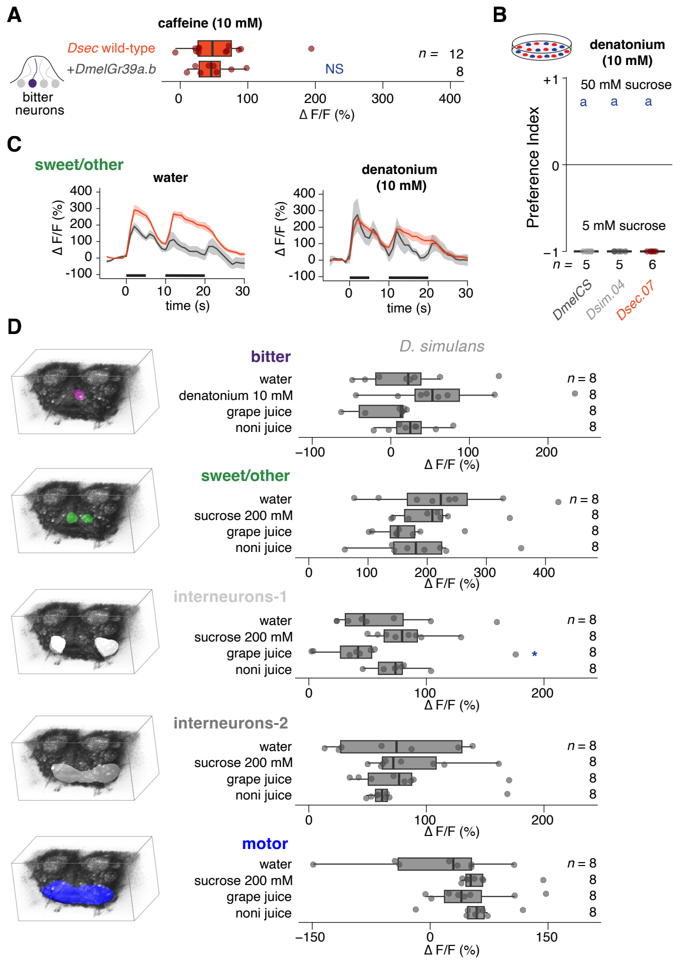
